# Optimizing the fertilizer N rates at different irrigation levels for optimum yield of wheat and corn at reduced nitrate leaching losses

**DOI:** 10.64898/2026.08.12.744458

**Authors:** Muhammad Tahir, David Mulla, Muhammad Zain, Saliha Maqbool, Anwar Ul Hassan, Muhammad Adeel

## Abstract

Optimum irrigation and fertilizer nitrogen (N) rates are important to improve crop yield at reduced environmental risks in the form of NO3-N leaching losses, without any financial loss. The study aimed to investigate the impact of rational irrigation and nitrogen management on wheat and maize crop yield vs. NO3-N leaching losses, with field experiments conducted at the experimental station, University of Agriculture Faisalabad, Pakistan, for two years, with wheat-fallow-corn seasons each year. Suction lysimeter were installed for collection of leachates while soil water balance was computed using the HYDRUS-1D model. We explored the various management strategies, including three irrigation and N levels (sub-optimal, optimal, and supra-optimal, referred to as I1, I2, and I3 for irrigation, and N1, N2, and N3 for nitrogen, respectively) for wheat and maize crops. The three irrigation levels were 325, 400, and 475 mm for wheat, and 375, 525, and 675 mm for maize crops. The three N-levels were 100, 130, and 160 kg ha-1 for wheat, and 220, 270, and 320 kg ha-1 for maize. The results indicated that increasing the irrigation and nitrogen levels significantly improved the growth and yield of both crops during both seasons. The highest grain yield of wheat (4.0 t ha-1) and maize (7.8 t ha-1) was observed with I3N3; however, I2N3 showed statistically no difference in yield, while showing significantly reduced (28.6%) annual NO3-N leaching losses of 23.8 kg ha-1, and the highest financial benefits of 780$. Sub-optimal levels of irrigation and N, though reduced the NO3-N leaching losses, caused significant yield losses, generally unacceptable to the farmers. Irrigation water use efficiency (WUEi) also improved by 12% in wheat and 20% in maize under I2 than that of the I3 level. Besides, considering economic profit, the highest value cost ratio (1.64 and 2.04 in wheat and maize, respectively) was achieved under the I2N3 treatment, as opposed to the other treatments. Based on comprehensive analysis, the I2N3 treatment is recommended for sustainable yield and minimal environmental risk in the wheat-maize cropping system. Moreover, it was observed that the rainy fallow period contributes 14.0-31.5% of the total NO3-N leaching losses. Further investigation is needed to minimize NO3-N leaching losses, especially during the rainy fallow period, by early maize sowing and increasing the efficiency of N fertilizer (such as fertilizer coating) under the flood irrigation system, to achieve the potential goals of sustainable productivity and environmental security.

## 1. Introduction

The rice-wheat cropping system is one of the largest agricultural production systems, covering about 13.5 million hectares in Asia, with 57% of it cultivated in South Asia [1,2]. This system provides the staple food to more than 20% of the South Asian and Chinese population [3]. However, continuous production of rice and wheat has resulted in different social, edaphic, and environmental issues. Thus, the productivity of this agricultural system is decreasing due to increased energy costs, depletion of groundwater resources, imbalanced soil fertility, and poor management of crop residues [4–6]. Thus, this production system forced the farmer community to shift towards alternative crops that require less water [7].

Maize is an important cereal crop that can be adapted under climatic variations [8,9]. Generally, in Pakistan, maize production was 9.8 million tons during the 2023 growing season, and it ranks fourth in terms of cultivated area after wheat, rice, and cotton [10]. Besides, the demand for maize is continuously increasing due to its huge nutritional benefits for the livestock industry. Further, the maize crop has a share of 2.9% in agricultural value addition and 0.7% in total GDP [10]. In many countries like Mexico, India, China, and Pakistan, the maize crop is preceded by wheat. The irrigated wheat-maize system is an important cropping system that covers nearly 6.1 million hectares in the western parts of Pakistan and India. Wheat crop is the most important crop in Pakistan, with production of 31.4 million tons during the 2023-24 growing season, and it is worth noting that wheat has a 2.2% share in GDP and a 9.0% share in the agricultural sector [10].

In Pakistan, many factors contribute to the low average yields of wheat and maize. Irrigation scheduling of field crops still disregards the soil-specific water requirements of crops and basic principles of sustainability and resource conservation. On the other hand, native fertility of agricultural soils in Pakistan is too low to support crop production, i.e., soil organic matter <1%, and is poor in nitrogen content, which cannot support sustainable agriculture [11]. However, intensive agriculture through irrigation and fertilizer management practices aimed at increasing crop yields introduced an enduring menace of groundwater pollution by unused fertilizers, such as nitrogen leaching from the irrigated fields. Moreover, flood irrigation applications or rainfall events that are out of phase with crop water uptake rates allow this additional water and soil solutes to be leached below the active root zone. Recently, groundwater pollution due to nitrate has been reported especially in the Punjab and Balochistan regions [12].

Unlike most of the developing countries, Pakistan consumes up to 83% of its groundwater resources for agricultural uses. Pakistan is deprived of proper management due to improper irrigation scheduling and application of outdated technologies, causing low WUE and crop yield [13]. Meanwhile, total surface available water in Pakistan has decreased by 10.6% over the last nine years from an initial value of 103.5 million acre-feet observed during 2015-2016 [10]. Due to water scarcity, increasingly degrading quality, and increased competition between water users, irrigated agriculture is focused on meeting water demands using new approaches in an environmentally friendly manner. Thus, the understanding of water dynamics of the soil profile of a crop under different water regimes is very important in determining the optimum irrigation scheduling for a crop in a specific area [14].

In Pakistan, due to inherently low organic carbon and poor nitrogen contents of agricultural soils, heavy application rates of nitrogen are usually applied, ranging between 100-200 and 200-350 kg ha-1 to support productive agriculture in wheat and maize crops, respectively. Thus, nitrogen applications should be optimized with irrigation events to ensure nitrogen availability to plants, because excessive nitrogen use not only leads to unfavorable increases in yield, but also results in environmental degradation such as soil acidification and greenhouse gas emissions [15–17]. Considering the above-mentioned issue, studying NO3- (end product of mineralization) leaching below the root zone is very important. Measuring nitrate leaching involves determining the water flux through a soil profile below the root zone over a known period and the averaged nitrate concentration. Furthermore, the application of simulation models to study the water dynamics of the soil profile of growing crops in the fields has been in practice [18]. Hydrus-1D model [19] is also widely used to simulate the one-dimensional movement of water in variably saturated soil.

Previously, studies focused on the effects of irrigation and nitrogen application on a single crop. However, to our best knowledge, a comprehensive study on wheat-maize productivity under different irrigation and nitrogen application levels has not been conducted for two consecutive growing seasons. Thus, a study was conducted in Pakistan region with objectives to (1) develop rational irrigation scheduling and nitrogen application level to obtain sustainable yield in wheat-maize cropping system by applying resources ranging from deficit to over, (2) assess the NO3-N leaching losses under different nitrogen rates receiving varying irrigation levels in wheat and maize crop, (3) recommend the efficient resource utilization strategy to farmers and stakeholders with aims to have minimum impacts on environment and agricultural system.

## 2. Materials and Methods

### 2.1. Description of experimental site

Field experiments were conducted at the Research Farm of the University of Agriculture, Faisalabad (31°26’ N and 73°06’ E), Punjab, Pakistan during 2007-08 and 2008-09. Wheat and maize crops were cultivated in a pattern of wheat-fallow-maize rotation. The experimental site has a semiarid climate, characterized by very hot and humid summers and dry, cool winters. The monthly meteorological data of precipitation, humidity, evaporation, mean maximum, and mean minimum temperatures of the experimental site from the local meteorological station are shown in Fig. 1. The experimental site received 517.1 mm and 373.4 mm precipitation during the years 2007-08 and 2008-09, respectively. However, the fallow period under wheat-fallow-maize rotation received 71.8 and 61.2% of precipitation during the respective years. The soil of the experimental site was classified as well-drained Hafizabad loam, mixed, semi-active, isohyperthermic Typic Calciargids.

**Fig. 1.**
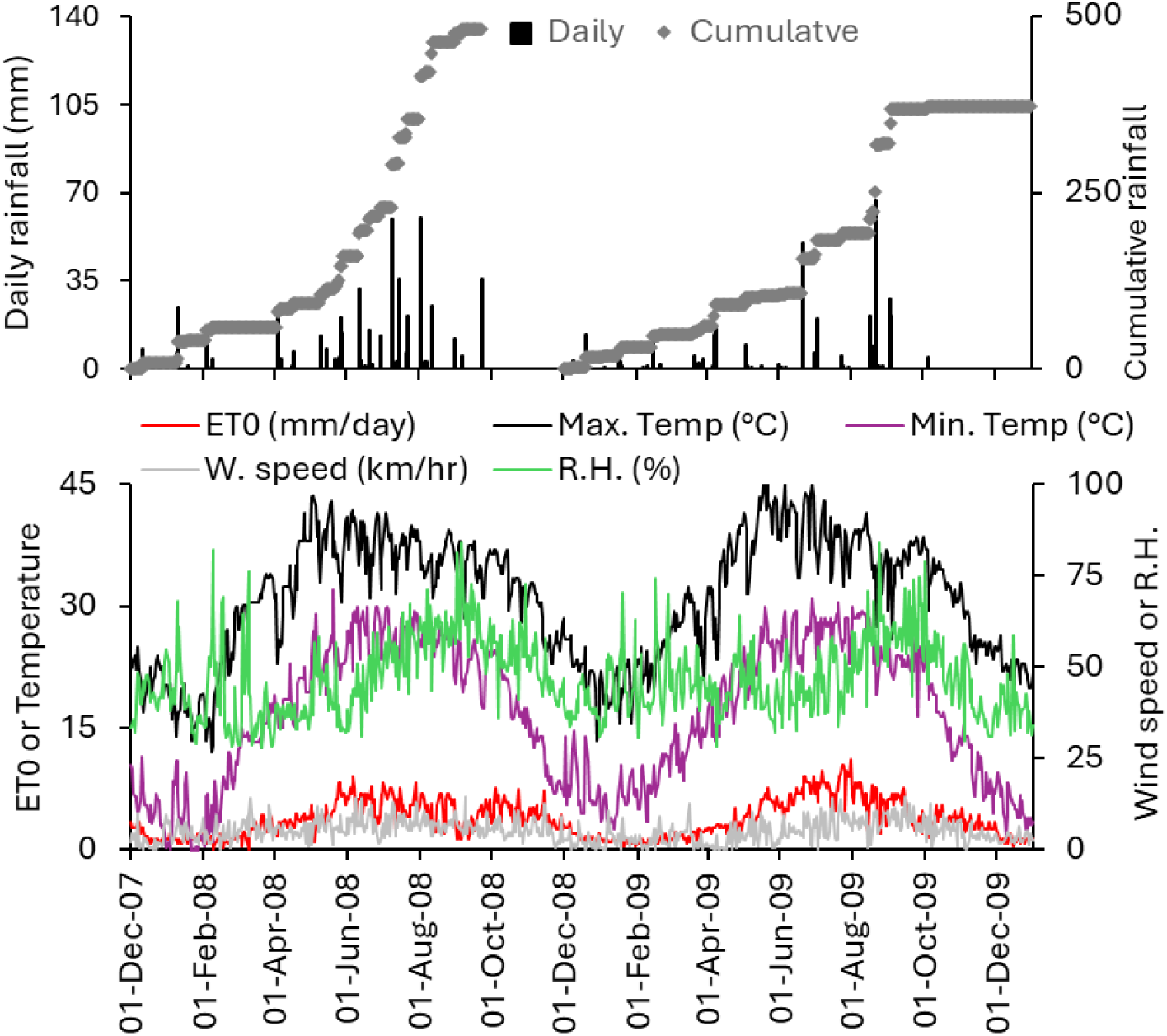
Precipitation and climatic conditions during the study period.

The physicochemical properties of the soil of the experimental site are shown in Table 1. Soil of the experimental site (Table 1) is poor in soil organic carbon (SOC), and calcareous in nature with a pH of 8.1. Soil organic carbon contents were observed 0.36, 0.25, and 0.19% at 0-0.35, 0.35-70 and 0.7-1.1 m depth, respectively. Bulk density (B.D.) of the soil ranged from 1.52-1.55 Mg m-3, where higher values observed at lower depths. Soil water contents at saturation (θs), field capacity (θ_FC_), and permanent wilting point (θ_PWP_) at different depths ranged from 0.41-0.43, 0.25-0.27, 0.11-0.12 cm^3^ cm^−3^, respectively. Soil saturated hydraulic conductivity (kfs) was observed to be higher (29.0 cm day^−1^) for 0-0.35 m depth, compared to that at 0.35-0.70 m (21.6 cm day^−1^) and 0.7-1.1 m depth (22.0 cm day^−1^).

**Table 1.** Measured soil physical and hydraulic parameters in the three main layers of the experimental site.

| Depth<br>(m) | Particle fraction (%) | | | B.D.<br>(Mg m<sup>-3</sup>) | $\theta_s$<br>-----cm<sup>3</sup> cm<sup>-3</sup>----- | $\theta_{FC}$<br>cm<sup>3</sup> cm<sup>-3</sup>----- | $\theta_{PWP}$<br>----- | $K_{fs}$<br>cm day<sup>-1</sup> | SOC<br>(%) |
| --- | --- | --- | --- | --- | --- | --- | --- | --- | --- |
|  | sand | silt | clay* |  |  |  |  |  |  |
| 0-0.35 | 39.0± | 36.0± | 25.0± | 1.52± | 0.43± | 0.26± | 0.11± | 29.0± | 0.36± |
|  | 0.25* | 0.16 | 0.17 | 0.04 | 0.01 | 0.014 | 0.008 | 0.64 | 0.03 |
| 0.35-0.70 | 40.5± | 34.0± | 25.5± | 1.56± | 0.41± | 0.27± | 0.11± | 21.6± | 0.25± |
|  | 0.31 | 0.25 | 0.21 | 0.03 | 0.01 | 0.009 | 0.014 | 0.65 | 0.01 |
| 0.70-1.10 | 43.0± | 32.0± | 25.0± | 1.55± | 0.42± | 0.25± | 0.12± | 22.0± | 0.19± |
|  | 0.20 | 0.25 | 0.15 | 0.02 | 0.01 | 0.012 | 0.012 | 0.78 | 0.01 |

\* Texture was loam according to USDA, ¶ Mean± standard error (data are average of three repeats); B.D., bulk density; $\theta_s$ , soil water content at saturation; $\theta_{FC}$ , soil water content at field capacity; $\theta_{PWP}$ , soil water content at permanent wilting point; $K_{fs}$ , soil saturated hydraulic conductivity.

### 2.2. Experimental design and crop management

Experiments were laid out using a randomized complete block split-plot design with three replications. Main plots consisted of three irrigation levels: sub-optimal (I1), optimal (I2), and supraoptimal (I3), while subplots contained three nitrogen levels: sub-optimal (N1), optimal (N2), and supra-optimal (N3). These three irrigation and nitrogen levels were 325, 400, and 475 mm, and 100, 130, and 160 kg N ha-1 for wheat; and 375, 525, and 675 mm, and 220, 270, and 320 kg N ha^−1^ for maize, respectively. The source of nitrogen fertilizer was urea. Wheat and maize crops received 85 and 140 kg ha^−1^ phosphorus as P_2_O_5_ and 65 and 105 kg ha^−1^ potassium as K_2_O in each plot before planting. Wheat and maize seeds were planted using a hand drill and dibbler, respectively. The size of each subplot was 5 m × 15 m. The planting density was 1.5 million plants ha-1 and 85,000 plants ha^−1^ for wheat and maize, respectively. Crop management practices were carried out according to local recommendations to achieve the potential grain yield (nitrogen and irrigation levels were the limiting factors). Weeds, diseases, and insects were intensively controlled throughout the growing period. Detailed information about the sowing and harvesting dates, cultivars, and the amount and timing of irrigation application is summarized in Table S1.

### 2.3. Leachate collection, drainage measurement and estimation of nitrate leaching

Suction lysimeters, equipped with ceramic cups, were installed at 1.1 m in each plot. These ceramic cups were embedded in the soil after the sowing of the first wheat crop. The preparation and insertion of the suction lysimeters followed the method stated by Webster et al. [20]. All the lysimeters were pre-treated with 1 M HC1. Leachates were collected at predefined intervals (Table S3) by applying a suction of 0.6 to 0.7 bar using a suction pump. Samples were analyzed within 24 hours of collection, otherwise stored at −4 °C.

Daily drainage was modeled and used to calculate daily and seasonal leaching losses. Soil water inputs were from irrigation and rainfall, while outputs included crop evapotranspiration (ET_c_) and drainage. Daily ET_c_ was calculated by multiplying the reference evapotranspiration (ET_0_) by crop coefficients adjusted for local climate conditions (Table 3) and a soil-water/nutrient stress factor (K_s_). The K_s_ or evapotranspiration reduction factor (ET_red_) was determined from the relationship between relative evapotranspiration reduction and relative yield reduction, using the method outlined by Doorenbos and Kassam [21]. We assumed no soil water and nutrient stress at the highest irrigation and N fertilizer rate. The ET_0_ for a hypothetical crop was calculated using the Penman-Monteith FAO-56 Equation [22].

HYDRUS-1D was used to estimate the drainage at 1.1 m depth by computing 1-dimensional water flux with soil water movement represented by Richard’s equation [23]. Soil water flux was considered vertical only (upward or downward), with surface runoff losses and sub-surface lateral flow assumed to be zero (<2% slope). Field/laboratory measured soil water retention and hydraulic parameters of three different soil layers were used to calibrate the model (Table 1). Parameters of the water retention curve (Table S2), optimized using RETC-fit software applying dual porosity - fit of retention *(Durner* model), were used for drainage calculation using HYDRUS-1D. The model was calibrated by optimizing the soil hydraulic parameters and comparing simulated versus calculated ET_c_. Daily NO_3_-N leaching (kg N ha^−1^ day^−1^) was then obtained by multiplying soil water NO_3_-N concentration (mg N L^−1^) by the estimated drainage below 1.2 m during the time interval preceding leachate collection. Seasonal NO_3_-N leaching was the sum of the daily leaching losses for the entire crop growth period.

Soil nitrate status was also monitored at different timings. Soil samples from 0-0.3, 0.3-0.7 and 0.70-1.10 m depth were collected and analyzed for NO_3_-N contents (nitrate build-up/depletion) at 0-1.1 m depth. The NO_3_-N at predefined time intervals was calculated by subtracting the current NO3-N mass from the initial mass. The NO_3_-N mass (kg ha^−1^) to a specific depth of soil was calculated by multiplying NO_3_-N concentration (mg kg^−1^) by the soil mass of one hectare for the respective depth. Nitrate-N was analyzed by the chromotropic acid method [24,25]. For measurement of soil nitrate, 10 g of air-dried soil was used, while for lysimeter samples, NO3-N concentration (ppm) was measured by taking a 1 mL leachate sample, following the same procedure as for soil samples.

### 2.4. Total available water and threshold value for readily available water

Total available water (*TAW*) was determined before each irrigation to assess any stress relative to *RAW* using the following formula:

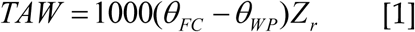

Where *TAW* is the total available water (mm), *θ_FC_* is the water content at field capacity (m^3^ m^−3^), *θ_WP_* is the water content at wilting point, and *Z_r_* is the rooting depth, which was taken as 1.1 m for both wheat and maize crops.

Readily available water (*RAW*) was calculated according to the following formula:

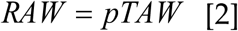

The value of p “threshold value for RAW” for wheat and maize (0.55) was taken from FAO Irrigation and Drainage Paper No. 33. and adjusted according to the following formula [22].

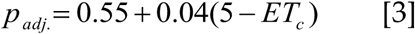

Where *p_adj_*is the adjusted fraction of *TAW* depleted from the root zone before any moisture stress and *ET_c_* as mm/day.

Critical point of *RAW* was calculated by the following equation:

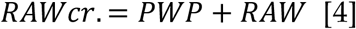

### 2.5. Plant parameters measurement

Root length density (RLD) and root weight density (RWD) of wheat and maize were measured at the completion of the mid-crop growth stage. Wheat roots were sampled from 0-15 cm depth within rows, using a root sampler (15 cm depth, 4 cm radius). Maize roots were sampled manually from a 0.35 m^3^ soil volume up to 35 cm depth. Roots were washed manually or by root washing systems, oven-dried at 65 °C until constant weight and then weighed [26]. RLD was measured using a *Dalta T-Scan* on a 5 g sub-sample. Leaf area index (LAI) was measured at 10 to 15 days’ intervals with a digital leaf area meter (LI-3000C) from 20 to 120 DAS and 10 to 115 DAS in wheat and maize, respectively. Plant height was measured using a meter tape throughout the growth period of wheat and maize crops. Water use efficiency (WUE) and irrigation water use efficiency (WUE_i_) were computed by dividing grain yield (kg ha-1) by ET_c_ (mm) and irrigation water use (mm), respectively.

### 2.6. Measurement of soil physicochemical properties

Oxidizable soil organic carbon (SOC), soil bulk density (B.D.), and the proportion of sand, silt, and clay were analyzed using the standard procedure given by Ryan et al. [24] (2001). Infiltration rate was measured with a double ring infiltrometer by taking readings at 5-minute intervals up to 30 minutes [27]. Soil saturated hydraulic conductivity (K_fs_) was measured by Guelph Permeameter (Model 2800 KI), taking three steady-state readings. The K_fs_ was then determined from the following formula:

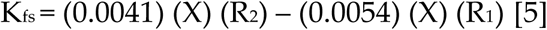

Where R_1_ and R_2_ are the steady-state rates of water-fall (cm s^−1^) in the reservoir at the first (*h_1_*) and second head (*h_2_*) of water (cm), respectively, and X (35.5 cm^2^) is the reservoir constant which is related to the cross-sectional area of the combined reservoir (cm^2^).

### 2.7. Statistical, economic, and marginal analysis

The data for crop yield, LAI, WUE, WUE_i_, plant height, RLD, RWD, deep drainage, nitrate-N leaching, and soil nitrate-N status under different irrigation and nitrogen co-limitation treatments were tested for homogeneity of variance using Levene’s test and for normality using the Shapiro-Wilks test before being subjected to analysis of variance (ANOVA) using GLM procedures in IBM SPSS Version 19.0 (IBM Corporation, Armonk, New York). The software packages STATISTIX 8.1 (StatSoft, Inc., 2001) and STATISTICA (Version 8, www.statsoft.com, OK 74104, US) were used for statistical analysis. The results showed a P value >0.05 for homogeneity of variance and <0.05 for the normality test, indicating that the data could be subjected to one-way ANOVA. The differences between treatment means were assessed using the least significant difference (LSD) test at a 95% level of significance. Data were analyzed for year effect using a two-tailed Student’s t-test. The effects of different irrigation and nitrogen treatments on wheat and maize crops were evaluated through economic and marginal analysis. Total input costs for each treatment, including variable and fixed costs, were calculated in US dollars after converting from Pakistani Rupees. Besides, a 10% downward yield adjustment was made to reflect the difference between the experimental yield and the expected yield on a farmer’s field. Net benefits for each treatment were calculated by subtracting total variable costs from total benefits. For Dominance analysis, treatments were given in order of increasing variable costs. A dominant (D) had net profit less than or equal to that of an experimental treatment with lower variable price. Marginal analysis was performed to determine the marginal rate of return (MRR) by dividing the change in cost and expressing it as a percentage [28]. Dominant treatments (D) were excluded from farmer recommendations due to their higher costs.

## 3. Results and Discussion

### 3.1. Effect of irrigation and nitrogen rates on growth characteristics of wheat and maize

Data presented in Fig. 2 represent the interactive effects of irrigation and nitrogen fertilizer treatments on plant height of wheat and maize during both study periods. The curves of plant height show that there were no significant differences in plant height during the early 60 and 40 days after sowing of wheat and maize, respectively, as the irrigation level did not change up to this timeframe. An increase in nitrogen fertilizer notably boosted plant height. Interestingly, a sharp rise in plant height after 50 and 40 days of sowing was observed in wheat and maize, respectively, reaching a maximum height at 90 and 60 days after sowing. Two-year average wheat plant height was highest by 15.16, 16.33, and 14.62% under I3N3 treatment at 70, 90, and 120 days after sowing than that of I1N1 treatment, respectively. The same trend was noticed in maize crop plant height, where I3N3 was higher by 10.18, 11.27, and 6.64% at 60, 70, and 100 days after sowing than I1N1.

**Fig. 2.**
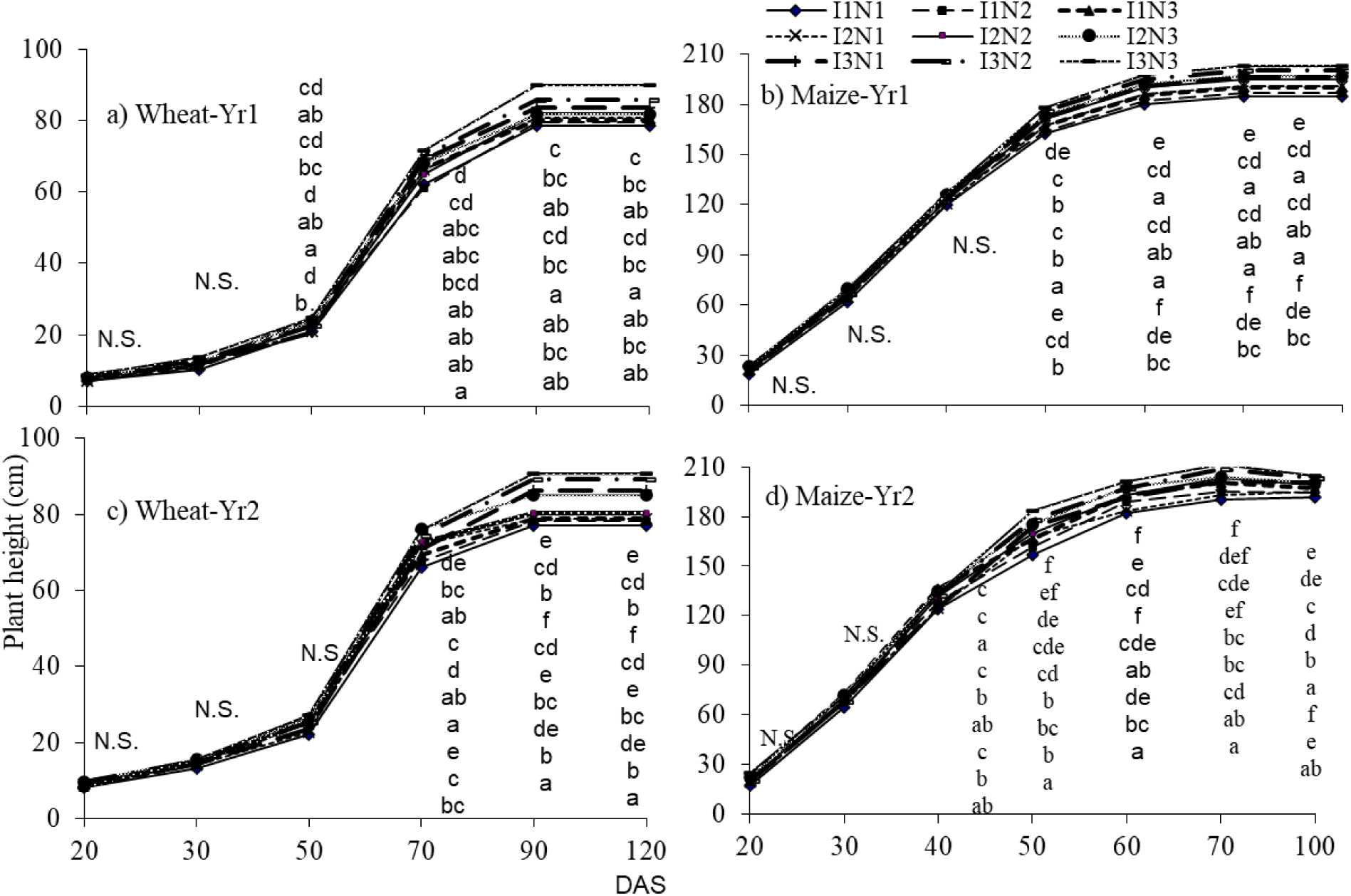
Effect of irrigation and nitrogen rates on plant height (cm) of wheat (a and c) and maize (b and d) during both growing seasons.

Results regarding the changes in LAI of wheat and maize crops due to various irrigation and nitrogen application rates are given in Fig. 3. We observed that LAI was non-significant in wheat and maize during the early days of crop growth in both study periods. As the crops continued to grow, a significant variation in LAI of both crops was observed. Similar to plant height, LAI also sharply increased at 50 DAS in wheat, reaching a maximum value of 4.67 and 4.93 at 80 DAS during Yr-1 and Yr-2, respectively, while a sharp increase at 40 DAS in the maize crop was observed, reaching a highest value of 5.47 and 5.40 at 80 DAS during respective years. We noticed that increasing the irrigation and nitrogen rates remarkably enhanced the LAI, as I3N3 increased the LAI by 12.98, 14.09, and 14.54% at 80, 100, and 110 DAS in the first wheat growing season. While the corresponding values during the second wheat growing season were 14.34, 15.00, and 13.38%, respectively. For maize, LAI in I3N3 was higher by 18.41, 14.61, and 17.17% at 80, 100, and 115 DAS than in I1N1 during the first growing season, while these values during the second growing season were 18.54, 26.76, and 27.11%, respectively.

**Fig. 3.**
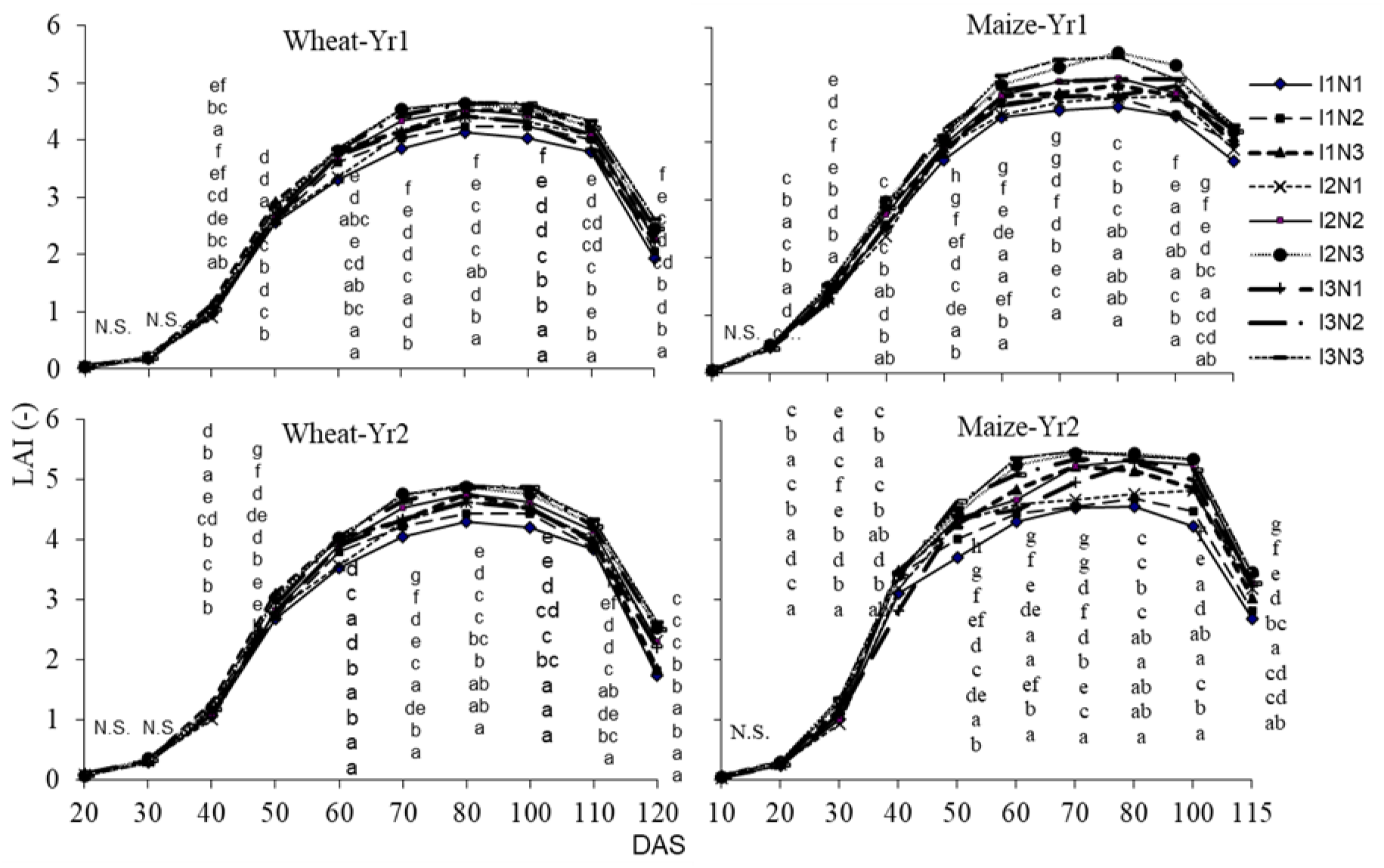
Effect of irrigation and nitrogen rates on LAI of wheat (a and c) and maize (b and d) during both growing seasons.

Increases in plant height and LAI due to high moisture and nitrogen fertilizer have been widely reported. Our results are consistent with previous studies [29,30], which found that increasing nitrogen availability enhances crop growth, while reduced irrigation decreases growth. Farooq et al. [31] reported that a decrease in irrigation and fertilizer reduced various cell processes such as cell elongation and cell division, ultimately leading to reduced plant height and LAI, as observed in the I1N1 treatment. Furthermore, previous studies [32] have demonstrated that higher nitrogen fertilizer application enhances the growth characteristics of wheat and maize. Thus, we conclude that the highest plant height and LAI under supra-optimal irrigation and nitrogen levels could be beneficial in terms of increasing the crop growth characteristics in both wheat and maize crops.

### 3.2. Effect of irrigation and nitrogen rates on root length density (cm cm-3) and root weight density (mg cm-3) of wheat and maize

Root development has a potential effect on crop water and nutrient uptake, which translates into high productivity. A mean comparison of root length density (RLD) and root weight density (RWD) under the interactive effect of irrigation and nitrogen fertilization treatments is given in Fig. 4. A significant difference among various treatments was observed in RLD and RWD of wheat and maize during both growing seasons. High irrigation and nitrogen applications remarkably enhanced RLD and RWD compared to low irrigation and nitrogen application treatments. The RLD of the wheat crop under I3N3 was 110 and 107.12% higher than the I1N1 treatment in the first and second growing seasons, respectively. Similarly, the RLD in maize was high by 142.62 and 49.31% in supra-optimal irrigation and nitrogen application (I3N3) as compared to the sub-optimal irrigation and nitrogen treatment (I1N1) during the first and second years, respectively.

**Fig. 4.**
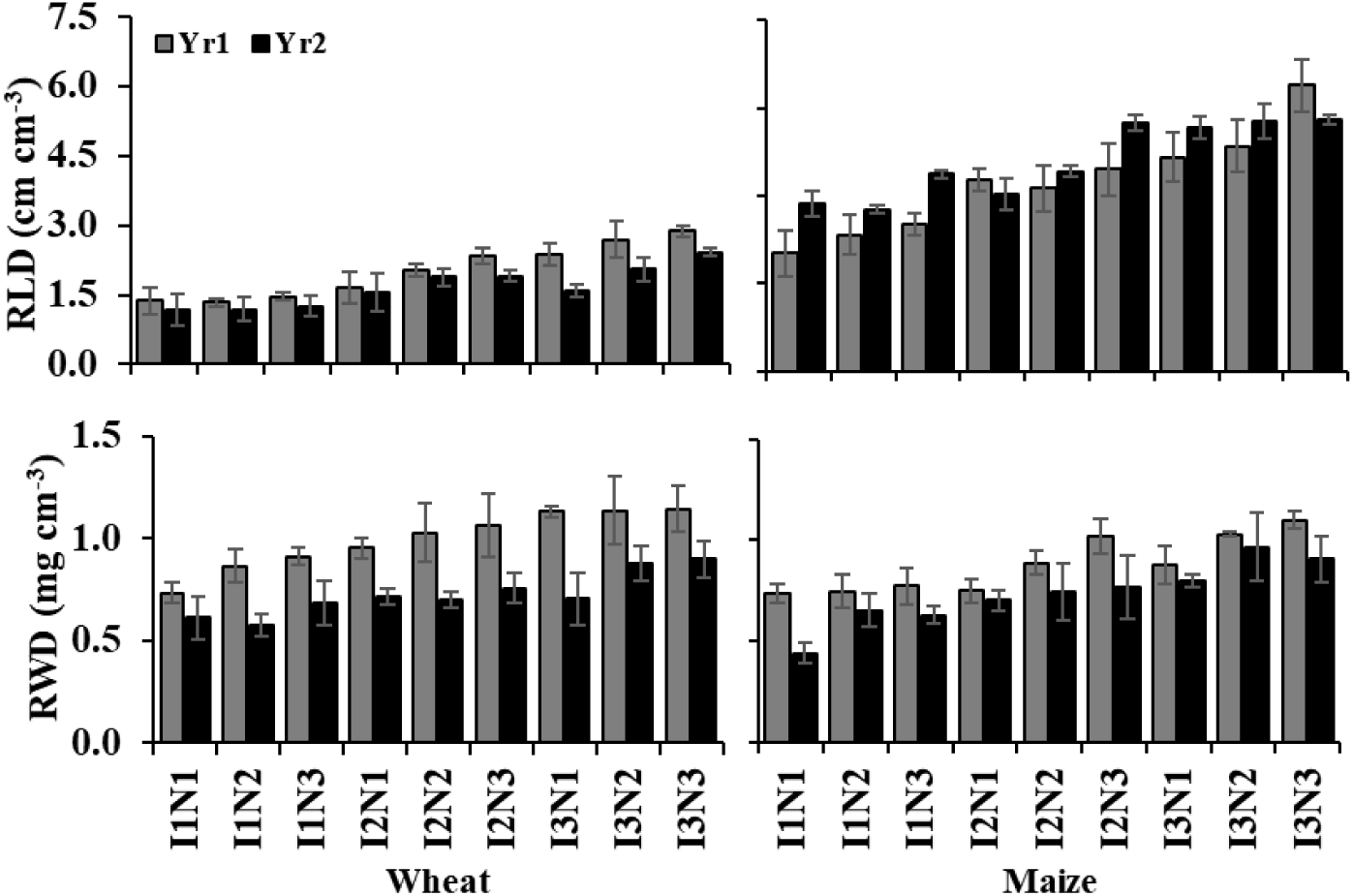
Effect of irrigation and nitrogen rates on root length density (cm cm^−3^) and root weight density (mg cm^−3^) of wheat and maize during both growing seasons.

Considering the effects of irrigation and nitrogen levels on RWD, the same trend was observed as in RLD during both study periods. We noticed that increasing the irrigation level from sub-optimal (I1) to supra-optimal (I3) increased the RWD by 36.08 and 32.79% in the first and second growing seasons of wheat, respectively. However, these values for the increase in maize RWD under I3 were 33.38 and 55.16% during the first and second experimental seasons, respectively. The RWD in the wheat crop was 55.91 and 47.30% higher under the I3N3 treatment as opposed to the I1N1 treatment in the first and second growing seasons, respectively. Similarly, the corresponding values of RWD for the maize crop were 49.32 and 106.06%, respectively, in both seasons. Interestingly, during the second year growing period, RLD of I3N2 was higher than I3N3 in the maize crop. The positive response of RLD was observed by Elazab et al. [33] and Man et al. [34] under high irrigation treatments. Our results are in line with Wang et al. [35], who found an increase in wheat growth under high irrigation compared with the no-irrigation treatment, and further stated that a decrease in nitrogen availability minimizes root growth. Furthermore, Buckley et al. [36] and Kang et al. [37] noticed that severe water stress disrupts the carbon and nitrogen balance, which leads to root cell death, damage to leaf tissues, and ultimately reduced photosynthesis. Similar results were reported in the maize crop under deficit irrigation treatment [38,39]. These results suggest that sub-optimal irrigation and nitrogen application or unmatched supply have unfavorable effects on root characteristics.

### 3.3. Length of growth stages and their Kc values for wheat and maize

The crop coefficient (Kc) values at different wheat and maize growth stages and the length of growth stages (days) are given in Table 2. The Kc values in the wheat crop range from 0.46 to 1.15 during both seasons, while for maize these values were between 0.50 and 1.18. For wheat, Kc values were 0.46, 1.15, 1.15, and 0.27 at initial, crop development, mid-stage, and late stage, respectively, during the first growing season. While the corresponding values during the second growing season were 0.47, 1.14, 1.14, and 0.44, respectively. Furthermore, the specific maize growth stage Kc values were 0.50, 1.17, 1.17, and 0.35 during the first study year, and were 0.51, 1.18, 1.18, and 0.58 during the second growing season at initial, crop development, mid-stage, and late-stage, respectively.

**Table 2.** Values of Kc for different growth stages (-) and their length (days).

| Crop | Growth stage |  |  |  |
| --- | --- | --- | --- | --- |
|  | Initial | Crop development | Mid stage | Late stage |
| Wheat-Yr1 | 0.46 (27±0.7*) | 0.46→1.15 (28±0.9) | 1.15 (60±1.4) | 1.15→0.27(26±0.6) |
| Maize -Yr1 | 0.50 (15±0.4) | 0.50→1.17 (28±1.4) | 1.17 (60±2.1) | 1.17→0.35(12±0.3) |
| Wheat -Yr2 | 0.47 (24±0.3) | 0.47→1.14 (32±0.6) | 1.14 (53±2.6) | 1.14→0.44(24±0.6) |
| Maize -Yr2 | 0.51 (14±1.1) | 0.51→1.18 (27±0.5) | 1.18 (61±1.5) | 1.18→0.58(10±1.1) |

\*Mean ± standard error (Values in the parentheses are the lengths of growth stages in days).

We found that the length of each growth stage in wheat and maize was slightly different during both growing seasons. The initial growth stage in wheat remained till 27 and 24 days during the first and second experimental periods. Similarly, the crop development, mid-stage, and late-stage lengths were 28, 60, and 26 days in the first wheat growing season, and 32, 53, and 24 days in the second growing season, respectively. Considering the maize crop, the length of initial, crop development, mid, and late stages was 15, 28, 60, and 12 days during the first growing season, and 14, 27, 61, and 10 days during the second growing season, respectively. The specific growth stage Kc values calculated in our study were smaller than those calculated by Li et al. [40] through the ratio of the measured ETc and the FAO-56 Penman-Monteith predicted ET0. According to Li et al. [40], the Kc values for wheat were 0.55, 1.03, 1.19, and 0.65 at initial, crop development, mid, and late stages, respectively, while for maize these values were 0.50, 1.02, 1.26, and 0.68, respectively. Besides, Kc values developed by Sharma et al. [41] were 0.36, 0.77, 1.05, and 0.25 at initial, development, mid, and end growth stages of the wheat crop, respectively. In another study, Liu et al. [42] also found slightly high Kc values (0.59, 1.24, 1.38, and 1.17) for the maize crop at the above-mentioned growth stages. This difference in Kc values during our experimental seasons might be attributable to the length of each growth stage.

### 3.4. Critical limit of readily available water and moisture content before each irrigation

The data regarding readily available water (θ_RAW_) and the moisture contents before irrigation are presented in Fig. 5. We found that the average values of θ_RAW_ were 0.176 and 0.173 during the first and second growing seasons of the wheat crop, respectively, while in the maize crop the average θ_RAW_ value was 0.178 during both growing seasons. Interestingly, the θ_RAW_ value in the wheat crop increases as the wheat continues to grow; however, in the maize crop, the θRAW value decreased from the initial to the end growth stage. This might be due to the differences in ETc during the crop growing season. In contradiction to our results, Wu et al. [43] measured slightly high θRAW values such as 0.244 cm^3^ cm^−3^ for loamy clay, 0.228 cm^3^ cm^−3^ for clay loam, and 0.190 cm^3^ cm^−3^ for sandy loam soil.

**Fig. 5.**
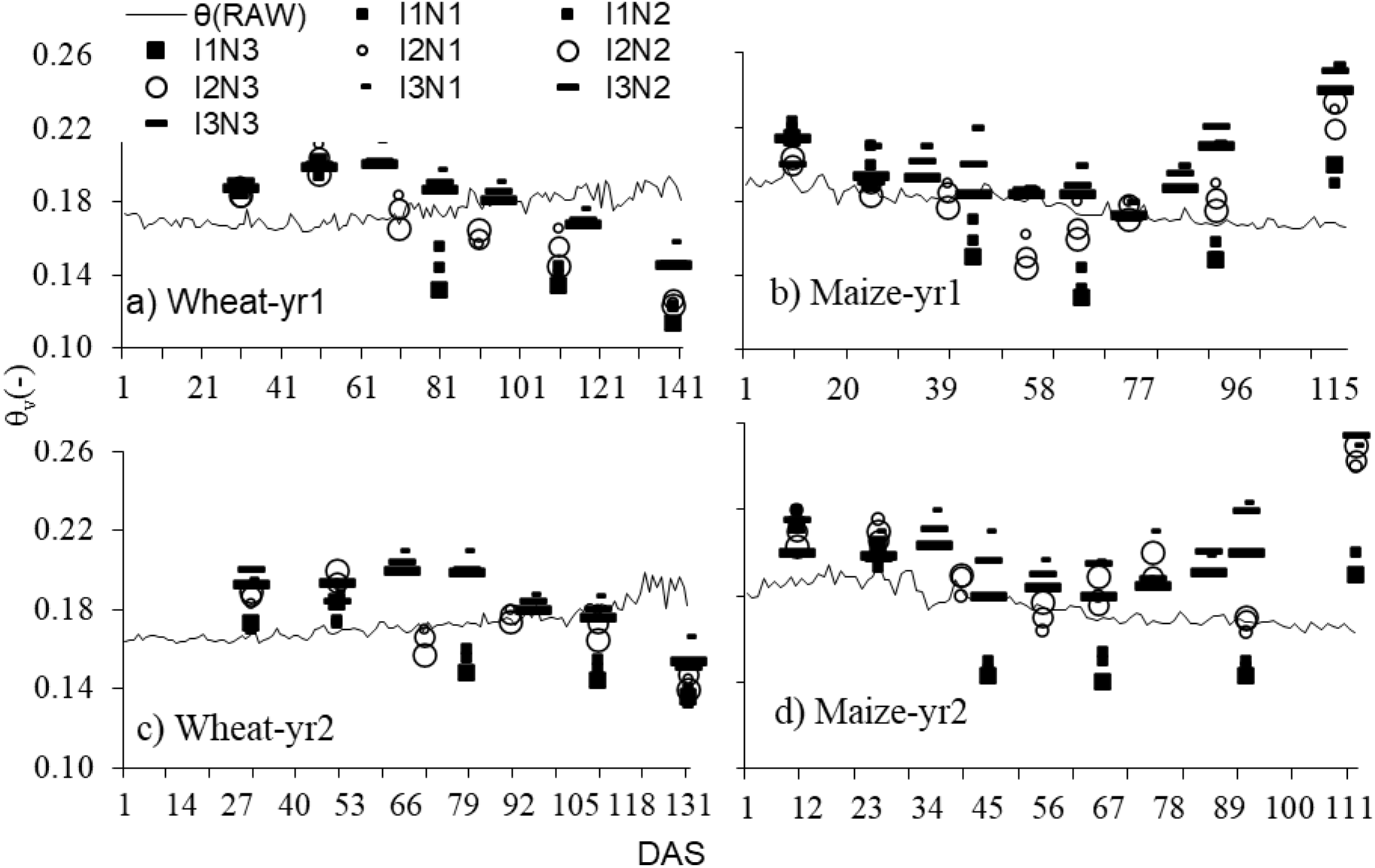
Critical limit of readily available water and moisture content before each irrigation.

A notable variation in available water content was observed in different irrigation and nitrogen fertilizer treatments. As the first and second irrigation times during wheat and maize were the same, the moisture contents were above θ_RAW_, and no stress was observed during this time span. Later, we found that the moisture contents at irrigation time were high in both maize and wheat under supraoptimal irrigation treatments (I3) with the same nitrogen application level, which indicates the high chances of drainage at later growth stages. However, the moisture contents in sub-optimal (I1) were below θRAW, indicating severe stress after the 2^nd^ irrigation. Considering the optimal irrigation level (I2) in the wheat crop, the moisture contents were below θ_RAW_ at the fourth and fifth irrigations in the first study period, representing a slight stress during this time, while there was no stress during the second growing season. Similar findings were also reported by He et al. [44], who found an increase in soil moisture contents with an increase in the irrigation level. This could also be due to increased soil salinity, as high salinity increases the readily available water for plants [45].

### 3.5. Effect of irrigation and nitrogen rates on grain yield (mg ha^−1^) and WUEi (kg ha^−1^ mm^−1^) of wheat and maize

The results regarding grain yield and irrigation water use efficiency of wheat and maize crops during both growing seasons are presented in Table 3. Different irrigation and nitrogen application treatments have significant effects on grain yield of wheat and maize, and a consistent response was found. It was observed that increasing the irrigation level from I1 to I3 increased the wheat grain yield by 26.26 and 27% during the first and second growing seasons, respectively. Similarly, increasing irrigation from super-optimal to supra-optimal in the maize crop enhanced grain yield by 40.40 and 50.87% during the first and second growing seasons, respectively. Furthermore, an increase in nitrogen level from N1 to N3 also significantly improved the grain yield by 17.02 and 20.89% in the wheat crop, while by 21.77 and 25% in the maize crop during the first and second study periods, respectively.

**Table 3.** Effect of irrigation and nitrogen rates on grain yield, WUEi of wheat and maize.

|  | Grain yield (Mg ha<sup>-1</sup>) |  |  |  | WUE<sub>i</sub> (kg ha<sup>-1</sup> mm<sup>-1</sup>) |  |  |  |
| --- | --- | --- | --- | --- | --- | --- | --- | --- |
|  | Wheat-1 | Wheat-2 | Maize-1 | Maize-2 | Wheat-1 | Wheat-2 | Maize-1 | Maize-2 |
| I<sub>1</sub>* | 3.58 b | 3.37 b | 6.04 b | 5.78 b | 0.90 a | 0.84 | 1.45 a | 1.39 ab |
| I<sub>2</sub> | 4.36 a | 4.19 a | 8.00 a | 8.30 a | 0.92 a | 0.88 | 1.42 a | 1.47 a |
| I<sub>3</sub> | 4.52 a | 4.28 a | 8.48 a | 8.72 a | 0.82 b | 0.78 | 1.18 b | 1.22 b |
| N<sub>1</sub> | 3.82 b | 3.59 b | 6.66 | 6.72 c | 0.81 b | 0.76 b | 1.19 b | 1.19 c |
| N<sub>2</sub> | 4.17 ab | 3.93 ab | 7.75 | 7.68 b | 0.88 a | 0.83 ab | 1.39 a | 1.37 b |
| N<sub>3</sub> | 4.47 a | 4.34 a | 8.11 | 8.40 a | 0.94 a | 0.92 a | 1.47 a | 1.52 a |
| I<sub>1</sub>N<sub>1</sub> | 3.39d° | 3.07 c | 5.13 e | 4.62 f | 0.85 bc | 0.77 ab | 1.23 cde | 1.11 d |
| - N<sub>2</sub> | 3.53 d | 3.32 bc | 6.28 de | 5.84 e | 0.88 abc | 0.83 ab | 1.51 abc | 1.40 bc |
| - N<sub>3</sub> | 3.82 cd | 3.73 a-c | 6.72 c-e | 6.89 d | 0.96 ab | 0.93 a | 1.62 a | 1.66 a |
| I<sub>2</sub>N<sub>1</sub> | 4.02 b-d | 3.71 a-c | 7.22 b-d | 7.60 cd | 0.85 c | 0.78 ab | 1.28 b-e | 1.34 cd |
| - N<sub>2</sub> | 4.40 a-c | 4.25 ab | 8.14 a-c | 8.23 bc | 0.93 abc | 0.90 ab | 1.44 a-d | 1.46 abc |
| - N<sub>3</sub> | 4.67 ab | 4.62 a | 8.63 ab | 9.06 a | 0.98 a | 0.97 ab | 1.53 ab | 1.60 ab |
| I<sub>3</sub>N<sub>1</sub> | 4.05 b-d | 3.98 a-c | 7.64 a-d | 7.93 c | 0.74 d | 0.72 b | 1.07 e | 1.11 d |
| N<sub>2</sub> | 4.58 ab | 4.21 ab | 8.82 a | 8.98 ab | 0.83 cd | 0.77 ab | 1.23 de | 1.26 cd |
| - N<sub>3</sub> | 4.92 a | 4.66 a | 8.97 a | 9.24 a | 0.89 abc | 0.85 ab | 1.25 b-e | 1.29 cd |
| LSD(p≤0.05) I | 0.27 | 0.61 | 0.78 | 1.47 | 0.06 | NS | 0.14 | 0.24 |
| N | 0.49 | 0.66 | 0.94 | 1.60 | 0.07 | 0.14 | 0.17 | 0.12 |
| I × N | 0.84,0.74 | 1.14,1.11 | 1.63,1.54 | 1.04,1.69 | 0.12, 0.11 | 0.25,0.24 | 0.29,0.28 | 0.21,0.29 |

\*\*I<sub>1</sub>, I<sub>2</sub>, and I<sub>3</sub> correspond to 325, 400, and 475 mm for wheat, and 375, 525, and 675 mm for the maize crop, respectively. \* N<sub>1</sub>, N<sub>2</sub>, and N<sub>3</sub> correspond to nitrogen 100, 130, and 160 kg ha<sup>-1</sup> for the wheat crop, and 220, 270, and 320 kg ha<sup>-1</sup> for the maize crop, respectively. \*Means sharing the same letter (s) do not differ significantly at $P < 0.05$ according to Duncan's Multiple Range Test. Year effect ( $T$ -test) was similar for grain yield, WUE<sub>i</sub> of wheat and maize.

Considering the interactive effect of irrigation and nitrogen fertilizer, the highest grain yield (4.92 and 4.66 mg ha^−1^ in wheat, 8.97 and 9.24 mg ha^−1^ in maize) was found in the I3N3 treatment during both study seasons. Further, the lowest grain yield (3.39 and 3.07 mg ha^−1^ in wheat, 5.13 and 4.62 mg ha^−1^ in maize) was noticed in I1N1 treatment in the first and second seasons of both crops, respectively. Interestingly, there was no statistical difference in grain yield of wheat and maize between I2N3 and I3N3 treatments in both experimental periods. Thus, we may suggest that the I2 irrigation level could be an acceptable treatment without significant loss in grain yield under the wheat-maize cropping system. Tahir et al. [46] found that a reduction in soil nitrogen by 7.31% and in nitrate concentration by 61.40% decreased the succeeding wheat crop production by 18.21%. Earlier studies also reported that grain yield is positively affected by the availability of nitrogen, while negatively affected by water stress [47,48], which is similar to our findings. Further, our results are in line with those of Si et al. [30], who reported that an increase in water and nitrogen significantly improved grain yield in wheat.

The variance analysis shows that WUEi was notably affected due to different irrigation and nitrogen treatments. It was noticed that WUEi values were high during the first growing season and were low during the second growing season of both crops. The increase in WUEi was observed with an increase in irrigation level from I1 to I2 (except during the first growing season of maize); however, I3 significantly reduced the WUEi in both seasons of wheat and maize. The WUEi increased by 12.20 and 12.82% in wheat, by 20.34 and 20.49% in I2, as compared to the I3 treatment in both seasons, respectively. In addition, increasing nitrogen application (N1 to N3) significantly increased WUEi in both crops. For example, the WUEi in N3 was higher than in N1 by 16.05 and 21.05% in wheat, and by 23.53 and 27.73% in maize during the first and second experimental periods, respectively. Considering the interactive effect of irrigation and nitrogen treatments, I2N3 showed the highest WUEi (0.98 and 0.97 kg ha-1 mm-1 in wheat, 1.53 and 1.60 kg ha-1 mm-1 in maize) compared with the other treatments during both seasons. However, the lowest values of WUEi (0.74 and 0.72 kg ha^−1^ mm^−1^ in wheat, 1.07 and 1.11 kg ha^−1^ mm^−1^ in maize) were noticed in the I3N1 treatment in the first and second growing periods, respectively. The high WUEi under I2N3 might be because of the greatest grain yield in this treatment as compared to other treatments. Our results corroborate Shen et al. [49], who reported that adding more irrigation water did not significantly improve water use efficiency. Zhao et al. [50] also reported that one-off irrigation in combination with adding nitrogen significantly enhanced water use efficiency compared with the other treatments. The increase in WUE (Table S4) was parallel to WUEi, indicating that N application also caused a significant increase in ET_c_ of both crops.

### 3.6. Effect of irrigation and nitrogen rates on evapotranspiration/evaporation (mm) during wheat-fallow-maize rotation

The variance analysis showed that crop evapotranspiration was remarkably affected due to various irrigation and nitrogen treatments, and their interactive effect was also significant during both growing seasons (Table 4). It was observed that increasing irrigation level notably enhanced the ETc in wheat and maize during both study periods, such as supra optimal irrigation level (I3) increased the ETc as compared to the I1 treatment by 18.86 and 16.84% in the first and second wheat seasons, while by 31.91 and 22.73% in the first and second maize seasons, respectively. Similarly, an increase in nitrogen level also enhanced the ETc. The average ETc values for nitrogen levels (N1, N2, and N3) were 400, 435, and 461 mm, respectively, in the wheat crop during the first year. While the corresponding values during the second study period were 397, 429, and 464 mm, respectively. Further, these values for the maize crop under various nitrogen levels (N1, N2, and N3) were 370, 430, and 450 mm in the first study period, and 358, 409, and 436 mm during the second growing season, respectively. Considering the interactive effects of irrigation and nitrogen fertilizer, we noticed that I3N3 has the highest ETc values, while irrigating with low irrigation and nitrogen levels (I1N1) significantly decreased the ETc in both crops during the whole experimental period. Moreover, I3N3 showed 33.24 and 36.39% more ETc than I1N1 in the first and second wheat growing seasons, respectively, while the corresponding values in maize were 63.33 and 47.77%, respectively.

**Table 4.**
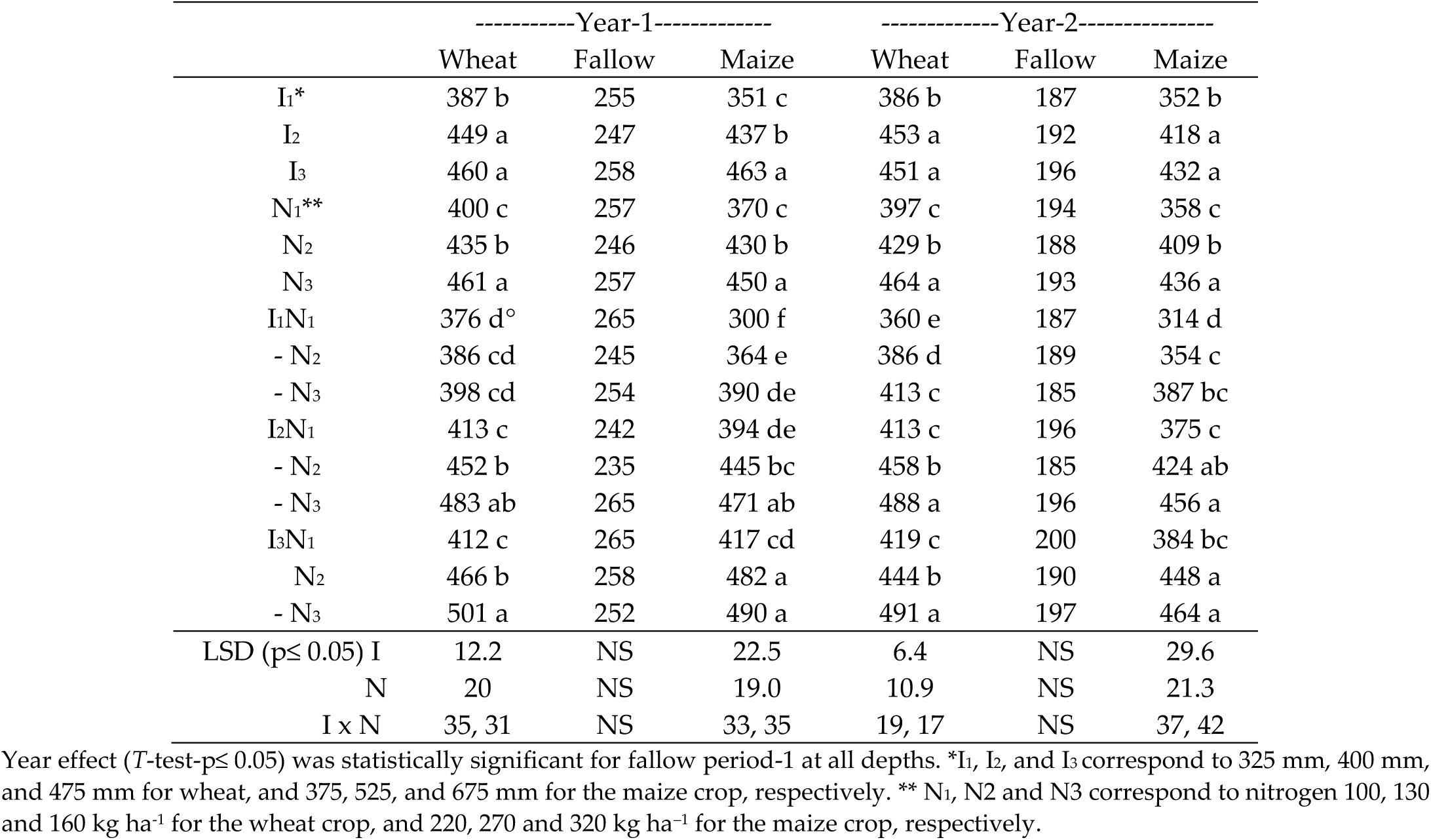
Effect of irrigation and nitrogen rates on evapotranspiration/evaporation (mm) during wheat-fallow-maize rotation.

|  | -----Year-1----- |  |  | -----Year-2----- |  |  |
| --- | --- | --- | --- | --- | --- | --- |
|  | Wheat | Fallow | Maize | Wheat | Fallow | Maize |
| I1* | 387 b | 255 | 351 c | 386 b | 187 | 352 b |
| I2 | 449 a | 247 | 437 b | 453 a | 192 | 418 a |
| I3 | 460 a | 258 | 463 a | 451 a | 196 | 432 a |
| N1** | 400 c | 257 | 370 c | 397 c | 194 | 358 c |
| N2 | 435 b | 246 | 430 b | 429 b | 188 | 409 b |
| N3 | 461 a | 257 | 450 a | 464 a | 193 | 436 a |
| I1N1 | 376 d° | 265 | 300 f | 360 e | 187 | 314 d |
| - N2 | 386 cd | 245 | 364 e | 386 d | 189 | 354 c |
| - N3 | 398 cd | 254 | 390 de | 413 c | 185 | 387 bc |
| I2N1 | 413 c | 242 | 394 de | 413 c | 196 | 375 c |
| - N2 | 452 b | 235 | 445 bc | 458 b | 185 | 424 ab |
| - N3 | 483 ab | 265 | 471 ab | 488 a | 196 | 456 a |
| I3N1 | 412 c | 265 | 417 cd | 419 c | 200 | 384 bc |
| N2 | 466 b | 258 | 482 a | 444 b | 190 | 448 a |
| - N3 | 501 a | 252 | 490 a | 491 a | 197 | 464 a |
| LSD (p≤ 0.05) I | 12.2 | NS | 22.5 | 6.4 | NS | 29.6 |
| N | 20 | NS | 19.0 | 10.9 | NS | 21.3 |
| I x N | 35, 31 | NS | 33, 35 | 19, 17 | NS | 37, 42 |

Year effect ( $T$ -test- $p \leq 0.05$ ) was statistically significant for fallow period-1 at all depths. \*I1, I2, and I3 correspond to 325 mm, 400 mm, and 475 mm for wheat, and 375, 525, and 675 mm for the maize crop, respectively. \*\* N1, N2 and N3 correspond to nitrogen 100, 130 and 160 kg ha<sup>-1</sup> for the wheat crop, and 220, 270 and 320 kg ha<sup>-1</sup> for the maize crop, respectively.

Data given in Table 4 show that evaporation measured in the fallow period was not significantly affected during both seasons. On average, evaporation under the I3 treatment was slightly higher than under the I2 and I1 treatments in both periods. Interestingly, it was noticed that overall evaporation during the first fallow period was 32.33% more than the second fallow period. This might be attributed to the more water input into the soil in the first growing season of the wheat and maize crops. It also indicates drought conditions in the second growing season and greater water demand. In both seasons, ETc increased with increasing irrigation and nitrogen, with the maximum value found in I3N3, which was similar to the findings of other studies [51]. The increase in evaporation under I3 may be attributed to high irrigation and ETc in this treatment. Srivastava et al. [52] also stated that maize evapotranspiration was high for irrigated and high nitrogen application treatments as compared to the no nitrogen treatment. Further, nitrogen application affects water consumption by improving soil water utilization [53].

### 3.7. Effect of irrigation and nitrogen rates on drainage (mm) at 110 cm depth during wheat-fallow-maize rotation

The results on Hydrus-1D simulated drainage are given for both growing seasons and the fallow period in Table 5. We found that drainage increased with increasing irrigation level; however, drainage decreased with increasing nitrogen level, as more water uptake occurs at high nitrogen levels. The drainage under N1 was greater than that under N3 by 45.10 and 37.25% in the first and second wheat seasons, respectively, while the corresponding values for the maize crop were 42.35 and 40.23%, respectively. Interestingly, during the fallow periods, I2 has the highest drainage, while I3 has the lowest drainage values. Drainage values were higher in the first growing season than in the second, which might be because of greater water input in the first season. Further, across all treatments, drainage was greater in the wheat crop than in the maize crop in both growing seasons. The interaction effect of irrigation and nitrogen application was also significant during both study periods. The highest values for drainage (187 and 180 mm in wheat, 204 and 192 mm in maize) were observed in the I3N1 treatment during the first and second study periods, respectively. The lowest values (68 and 81 mm in wheat, 37 and 39 mm in maize) were observed in the I1N3 treatment, respectively. The high drainage under the I3 treatment might occur through pass flow due to wetter soil [54]. However, the low drainage found in the N3 treatment might be due to dense crops or improved crop growth characteristics, which leads to greater consumption of irrigation water and ultimately low drainage [55,56].

**Table 5.** Effect of irrigation and nitrogen rates on drainage (mm) at 110 cm depth during wheat-fallow-maize rotation.

|  | Year-1 |  |  | Year-2 |  |  |
| --- | --- | --- | --- | --- | --- | --- |
|  | Wheat | Fallow | Maize | Wheat | Fallow | Maize |
| I1* | 83 c | 109 | 53 c | 120 b | 66 | 54 c |
| I2 | 131 b | 110 | 94 b | 118 ab | 68 | 90 b |
| I3 | 166 a | 97 | 169 a | 154 a | 64 | 165 a |
| N1** | 148 a | 114 a | 121 a | 140 a | 70 a | 122 a |
| N2 | 129 b | 100 b | 110 b | 117 b | 63 b | 101 b |
| N3 | 102 c | 106 ab | 85 c | 102 c | 64 b | 87 c |
| I1N1 | 93 fg | 109 bc | 54 f | 91 ef | 57 c | 69 f |
| - N2 | 88 g | 112 ab | 67 e | 87 fg | 69 a | 54 g |
| - N3 | 68 h | 102 bcd | 37 g | 81 g | 60 bc | 39 h |
| I2N1 | 165 d | 114 ab | 104 d | 152 bc | 68 ab | 105 d |
| - N2 | 133 e | 125 a | 97 d | 112 de | 73 a | 91 e |
| - N3 | 95 f | 109 bc | 80 e | 91 fg | 70 a | 76 f |
| I3N1 | 187 a | 97 cd | 204 a | 180 a | 60 bc | 192 a |
| N2 | 167 b | 104 bc | 166 b | 151 b | 68 ab | 158 b |
| - N3 | 144 c | 89 d | 137 c | 134 cd | 60 bc | 145 c |
| LSD (p≤ 0.05) I | 20.8 | NS | 12.9 | 20.9 | NS | 8.2 |
| N | 6.3 | 7.4 | 7.3 | 8.2 | 4.0 | 4.4 |
| I × N | 13, 22 | 13, 14 | 13, 16 | 14, 24 | 7, 9 | 8, 10 |

Year effect ( $T$ -test- $p \leq 0.05$ ) was statistically significant for fallow period-1 at all depths. \*I1, I2 and I3 correspond to 325 mm, 400 mm and 475 mm for wheat, and 375, 525 and 675 mm for maize crop, respectively. \*\* N1, N2 and N3 correspond to nitrogen 100, 130 and 160 kg ha<sup>-1</sup> for the wheat crop, and 220, 270 and 320 kg ha<sup>-1</sup> for the maize crop, respectively.

### 3.8. Effect of irrigation and N rates on NO_3_^−^-N leaching losses (kg ha^−1^) during wheat-fallow-maize rotation

The NO_3_-N leaching during the crop growth season and fallow period is given in Table 6. The value of NO_3_-N leaching during the first wheat period was in the range of 4.7-9.2 and 4.3-11.3 kg ha^−1^ during the first and second years. However, the corresponding values for the maize crop were 3.6-11.8 kg ha-1 in the first study period and were in the range of 4.3-14.0 kg ha-1 during the second study period. We found an increasing trend in NO_3_-N leaching with increased irrigation and nitrogen application during both growing seasons of wheat and maize. For example, the leaching losses under supra-optimal irrigation (I3) to wheat were 8.4 and 10.2 kg ha^−1^ in the first and second growing seasons, respectively, indicating about 64.71 and 108.16% higher losses compared to sub-optimal irrigation (I1) in both seasons, respectively. Similarly, the losses under I3 in maize growing seasons were 11.7 and 13.2 kg ha^−1^, which indicates 192.50 and 186.96% higher losses than the I1 treatment. Furthermore, increasing the nitrogen level from N1 to N3 increased the leaching losses by 16.13 and 22.39% in both seasons of the wheat crop, while by 1.43 and 7.23% in the first and second growing seasons of maize, respectively. Considering the interactive effect of irrigation and nitrogen, the highest losses (9.2 and 11.3 kg ha-1 in wheat) were observed in the I3N3 treatment during both seasons. However, in maize, the losses were more in the I3N2 (11.8 kg ha-1) and I3N3 (14 kg ha-1) treatments in the first and second growing seasons, respectively. Furthermore, the lowest losses were noticed in the I1N1 treatment during both growing seasons of wheat and maize crops. In addition, irrigation had non-significant effects on NO_3_-N leaching during both fallow periods.

**Table 6.** Effect of irrigation and nitrogen rates on NO_3_-N leaching losses at 110 cm depth during wheat-fallow-maize rotation.

|  | Year-1 |  |  | Year-2 |  |  |
| --- | --- | --- | --- | --- | --- | --- |
|  | Wheat | Fallow | Maize | Wheat | Fallow | Maize |
| I<sub>1</sub>* | 5.1 c | 4.7 | 4.0 c | 4.9 c | 3.3 | 4.6 c |
| I<sub>2</sub> | 6.3 b | 5.5 | 6.1 b | 8.0 b | 3.3 | 7.9 b |
| I<sub>3</sub> | 8.4 a | 4.5 | 11.7 a | 10.2 a | 3.3 | 13.2 a |
| N<sub>1</sub>** | 6.2 b | 4.8 ab | 7.0 | 6.7 b | 3.0 | 8.3 |
| N<sub>2</sub> | 6.4 b | 5.3 a | 7.5 | 8.1 a | 3.6 | 8.5 |
| N<sub>3</sub> | 7.2 a | 4.6 b | 7.1 | 8.2 a | 3.4 | 8.9 |
| I<sub>1</sub>N<sub>1</sub> | 4.7 e | 4.7 b | 3.6 d | 4.3 | 3.1 | 4.5 c |
| - N<sub>2</sub> | 4.9 de | 4.9 b | 4.4 cd | 5.3 | 3.6 | 4.9 c |
| - N<sub>3</sub> | 5.7 cd | 4.5 b | 3.9 d | 5.1 | 3.2 | 4.3 c |
| I<sub>2</sub>N<sub>1</sub> | 6.2 c | 5.1ab | 5.7 bc | 7.4 | 3.1 | 7.6 b |
| - N<sub>2</sub> | 6.3 c | 6.3 a | 6.4 b | 8.2 | 3.4 | 7.9 b |
| - N<sub>3</sub> | 6.5 c | 5.1 ab | 6.1 bc | 8.3 | 3.6 | 8.3 b |
| I<sub>3</sub>N<sub>1</sub> | 7.8 b | 4.7 b | 11.8 a | 8.4 | 2.8 | 12.8 a |
| N<sub>2</sub> | 8.1 b | 4.7 b | 11.8 a | 10.9 | 3.7 | 12.7 a |
| - N<sub>3</sub> | 9.2 a | 4.1 b | 11.4 a | 11.3 | 3.4 | 14.0 a |
| LSD (p≤ 0.05) I | 0.98 | NS | 1.19 | 2.17 | NS | 2.22 |
| N | 0.54 | 0.24 | NS | 1.03 | NS | NS |
| I × N | 0.94, 1.23 | 1.25, 1.04 | 1.42, 1.65 | NS | NS | 1.91, 2.69 |

Year effects ( $T$ -test $p \leq 0.05$ ) were statistically significant for fallow period and maize. \*I<sub>1</sub>, I<sub>2</sub> and I<sub>3</sub> correspond to 325 mm, 400 mm and 475 mm for wheat, and 375, 525 and 675 mm for maize crop, respectively. \*\* N<sub>1</sub>, N<sub>2</sub> and N<sub>3</sub> correspond to nitrogen 100, 130 and 160 kg ha<sup>-1</sup> for wheat crop, and 220, 270 and 320 kg ha<sup>-1</sup> for maize crop, respectively.

The results showed that a reasonable decrease in irrigation and nitrogen levels can remarkably reduce the NO_3_-N leaching losses. Similar to the current study, Souza et al. [18] also determined nitrate leaching by calculating water drainage through Hydrus-1D and found that nitrogen treatments have significant effects on nitrate leaching. Our results are in line with the earlier studies of Li et al. [57] and Lu et al. [58], who reported increased inorganic nitrogen leaching losses with an increase in irrigation depth. Furthermore, an increase in nitrogen application may increase the available nitrogen in the root zone and also the NO_3_-N leaching losses [59,60].

### 3.9. Effect of irrigation and N rates on NO_3_-N built-up (kg ha^−1^) in wheat-maize cropping system

The soil nitrate nitrogen status was measured at 60 DAS, harvest, and fallow period in both growing seasons of wheat and maize, and the results are presented in Table 7. We noticed that increasing irrigation level significantly reduced the soil nitrogen in wheat and maize after 60 days of sowing in both study periods. For instance, across the nitrogen levels, the I1 treatment had 27.89% and 8.89% more nitrogen than the I3 treatment at 60 DAS in the first and second growing seasons of wheat, respectively, while these values after 60 DAS in the maize crop were 25.64 and 13.51%, respectively. Interestingly, soil nitrate nitrogen contents were high at harvest and the fallow period under the I3 treatment as compared to the I1 treatment. Regardless of irrigation, increasing the nitrogen level notably enhanced the NO_3_-N contents in soil at all observation times (60 DAS, harvest, fallow) in the wheat-maize cropping system. Taking into consideration the interactive effect (I × N), the NO3-N contents ranged between 11.3-37.3 kg ha-1 in the first wheat-maize cropping system, and between 7.8-33.4 kg ha-1 during the second growing season of wheat and maize. At 60 DAS, the highest soil inorganic nitrogen was noticed in I1N3 (25.4 and 19.4 kg ha^−1^ in wheat, 37.3 and 33.4 kg ha^−1^ in maize), which was at par with the I2N3 treatment during both study seasons of wheat and maize, respectively. However, the lowest soil nitrogen values at 60 DAS were observed in the I3N1 treatment (11.3 and 7.8 kg ha^−1^ in wheat, 19.3 and 18.7 kg ha^−1^ in maize) in both seasons, respectively. It was observed that the lowest soil nitrogen values were in I2N1 after both wheat harvest seasons, and in the I3N1 treatment after maize harvest during both seasons. On the other hand, the highest soil nitrogen values were not in the same treatment after wheat and maize harvest. For example, the highest values (7.6 and 6.7 kg ha^−1^) after wheat harvest were noticed in I3N3 and I2N3 after the first and second seasons of wheat harvest, respectively, while the highest values (11.2 and 6.4 kg ha-1) after maize crop harvest were recorded in I1N3 and I3N3 treatments, respectively. This might be due to more precipitation during the first growing season than during the second. The nitrate nitrogen contents in soil were lower during both fallow periods because these periods received about 71.8 and 61.2% of the total precipitation in the first and second growing seasons, respectively, which might lead to more leaching of nitrogen in deeper soil layers or might cause soil erosion. Tahir et al. [61] stated that heavy rainfall in the monsoon season, with peaks occurring specifically in April and June, may cause soil erosion. Our results are consistent with previous findings that high nitrogen application markedly increases soil available nitrogen [44,62]. Further, based on our findings, we suggest that optimizing irrigation and nitrogen application is a potential strategy for reducing nitrate nitrogen residues and improving their distribution in the soil profile [63–65].

**Table 7.** Effect of irrigation and nitrogen rates on NO_3_^−^-N built-up (kg ha^−1^) during wheat, fallow period, and maize at 0-110 cm depths.

|  | -----Year-1----- |  |  |  |  | -----Year-2----- |  |  |  |  |
| --- | --- | --- | --- | --- | --- | --- | --- | --- | --- | --- |
|  | Wheat |  | Fallow | Maize |  | Wheat |  | Fallow | Maize |  |
|  | 60d | Harv. | end | 60d | Harv. | 60d | Harv. | end | 60d | Harv |
| I<sub>1</sub>* | 18.8 a | 2.9 b | -0.5 | 29.4 b | 7.7 b | 14.7 a | 4.7 b | -4.0 b | 25.2 | 3.7 |
| I<sub>2</sub> | 16.3 ab | 3.5 ab | -0.6 | 25.2 ab | 6.2 ab | 13.7 b | 5.3 ab | -5.1 b | 22.7 | 3.0 |
| I<sub>3</sub> | 14.7 b | 4.3 a | 0.5 | 23.4 a | 5.8 a | 13.5 b | 5.5 a | -1.2 a | 22.2 | 3.8 |
| N<sub>1</sub>** | 12.0 c | 0.8 c | -2.3 c | 20.0 c | 3.3 c | 9.0 c | 3.9 c | -4.7 c | 17.3 c | 1.3 c |
| N<sub>2</sub> | 15.2 b | 4.0 b | -0.1 b | 25.2 b | 7.0 b | 13.0 b | 5.4 b | -3.4 b | 22.4 b | 3.7 b |
| N<sub>3</sub> | 22.7 a | 5.8 a | 1.9 a | 32.7 a | 9.4 a | 20.0 a | 6.2 a | -2.2 a | 30.5 a | 5.4 a |
| I<sub>1</sub>N<sub>1</sub> | 12.3 de | 1.2 e | -1.4 cd | 21.5 de | 4.2 c | 10.3 e | 3.3 e | -5.3 fg | 17.8 ef | 2.3 d |
| - N<sub>2</sub> | 18.7 c | 3.2 d | -1.2 c | 29.3 bc | 7.7 b | 14.5 c | 5.5 b | -4.3 de | 24.5 cd | 4.3 c |
| - N<sub>3</sub> | 25.4 a | 4.3 c | 1.2 b | 37.3 a | 11.2 a | 19.4 ab | 5.4 b | -2.3 bc | 33.4 a | 4.4 c |
| I<sub>2</sub>N<sub>1</sub> | 12.3 de | 0.4 f | -3.2 e | 19.3 e | 3.2 c | 9.0 ef | 3.9 d | -5.4 g | 15.4 f | 1.2 e |
| - N<sub>2</sub> | 14.4 d | 4.4 c | 0.3 b | 24.7 cd | 6.5 b | 13.3 cd | 5.3 b | -4.5 ef | 22.4 de | 2.3 d |
| - N<sub>3</sub> | 22.3 b | 5.6 b | 1.2 b | 31.5 b | 8.8 b | 18.9 b | 6.7 a | -5.4 g | 30.4 ab | 5.4 b |
| I<sub>3</sub>N<sub>1</sub> | 11.3 e | 0.7 f | -2.3 de | 19.3 e | 2.5 c | 7.8 f | 4.6 c | -3.3 cd | 18.7 ef | 0.4 f |
| N<sub>2</sub> | 12.4 de | 4.5 c | 0.5 b | 21.7 de | 6.7 b | 11.2 de | 5.4 b | -1.4 b | 20.4de | 4.5 c |
| - N<sub>3</sub> | 20.3 bc | 7.6 a | 3.4 a | 29.3 bc | 8.3 b | 21.6 a | 6.5 a | 1.2 a | 27.6 bc | 6.4 a |
| LSD (p≤ 0.05) I | 1.36 | 0.30 | NS | 3.97 | 0.89 | 0.78 | 0.24 | 0.79 | NS | NS |
| N | 1.87 | 0.35 | 0.62 | 1.75 | 1.42 | 1.67 | 0.29 | 0.50 | 2.19 | 0.44 |
| I × N | 3.24, | 0.43, | 1.07, | 3.02, | 2.46, | 2.90, | 0.36, | 0.87, | 3.80, | 0.77, |
|  | 2.96 | 0.41 | 1.06 | 4.65 | 2.19 | 2.49 | 0.33 | 1.06 | 4.89 | 0.68 |

Note. Soil NO<sub>3</sub>-N built-up (0-110 cm) was measured at 60 days after sowing and at harvest of wheat and the maize crop. During the fallow period, it was measured at the end of the fallow period.

### 3.10. Effect of irrigation and nitrogen rates on N recovery (%) by wheat and maize

The nitrogen recovery (%) by wheat and maize crop was measured based on nitrogen inputs and outputs calculation as given in Table 8. Increasing the irrigation level from I1 to I3 increased the N recovery in wheat and maize during the first and second growing seasons. For example, N recovery under I3 irrigation was 75.6 and 71.6% in wheat, while it was 63.1 and 65.5% in maize during the first and second growing seasons, respectively. However, the corresponding values under I1 were 57.2 and 55.4% in wheat, while 51 and 53.4% in the maize crop during both study periods, respectively. On average across the three irrigation levels, increasing nitrogen application notably decreased the recovery percentage in the wheat-maize cropping system. The nitrogen recovery in the wheat crop under N1 was 73.7 and 73.3% as compared to 61.2 and 59.7% under N3 in the first and second growing seasons, respectively. However, in the maize crop, N recovery slightly improved (3.70 and 2.42% during the first and second years, respectively) by increasing N from N1 to N2, but it significantly decreased at the supra-optimal level. The highest nitrogen recovery in the maize crop was 61.7 and 63.6% under N2 treatment, while the lowest nitrogen recovery values were 54.9 and 59.3% in N3 during the first and second study periods, respectively. Considering the interactive effect of irrigation and nitrogen levels on nitrogen recovery percentage, the values ranged between 49.7 and 79.5% during the first growing season, and between 55.2 and 80.4% in the second experimental year. The nitrogen recovery values in wheat growing seasons were higher than those in the maize cropping system. We found that I3N1 has the highest recovery values (79.5 and 80.4% in wheat, 66.3 and 66.1% in maize) as compared to the other treatments during the first and second study periods, respectively. However, the lowest values for nitrogen recovery were identified under different treatments in the wheat-maize cropping system. Such as, I1N3 had the lowest values (49.7 and 55.2%) in the wheat crop growing seasons, respectively, while in the maize crop, the minimum nitrogen recovery values (48.4 and 48.7% during the first and second years, respectively) were observed in I1N1 compared to those of other treatments. An adequate supply of nitrogen is critical for crop growth and development [66]. Based on our results, we found that nitrogen application under N1 and N2 treatments could be the potential rates for wheat and maize, respectively. However, high nitrogen application (N3) could not be a feasible rate under the wheat-maize cropping system. Further, our results are similar to previous studies, which have shown that reasonable nitrogen rates, combined with irrigation levels, promote nitrogen absorption, leading to increased irrigation and nitrogen efficiency and ultimately improved grain yield [67–69].

**Table 8.** Effect of irrigation and nitrogen rates on N recovery (%) by wheat and maize.

|  | Year-1<br>Wheat | Year-1<br>Maize | Year-2<br>Wheat | Year-2<br>Maize |
| --- | --- | --- | --- | --- |
| I1* | 57.2c | 51.0 b | 55.4b | 53.4b |
| I2 | 70.9b | 62.0 a | 70.2a | 66.1a |
| I3 | 75.6a | 63.1 a | 71.6a | 65.5a |
| N1** | 73.7a | 59.5ab | 73.3a | 62.1 |
| N2 | 68.2a | 61.7a | 64.1b | 63.6 |
| N3 | 61.2b | 54.9b | 59.7b | 59.3 |
| I1N1 | 63.7cd° | 48.4c | 61.2cd | 48.7d |
| -N2 | 58.2de | 54.8abc | 52.6d | 57.4bcd |
| -N3 | 49.7e | 49.9bc | 52.2d | 54.2cd |
| I2N1 | 77.8ab | 63.8ab | 77.2ab | 71.4a |
| -N2 | 69.7bc | 64.3ab | 69.6abc | 64.7abc |
| -N3 | 65.3cd | 58.0abc | 63.7bcd | 62.0abc |
| I3N1 | 79.5a | 66.3a | 80.4a | 66.1ab |
| -N2 | 76.6ab | 66.0a | 70.0abc | 68.8ab |
| -N3 | 68.6bc | 57.0abc | 63.1bcd | 61.5abc |
| LSD* (p≤ 0.05) I | 4.5 | 7.6 | 9.4 | 8.5 |
| N | 6.3 | 6.1 | 7.9 | NS |
| I × N | 10.9,9.9 | 10.5,15.2 | 13.7, | 9.6,11.5 |

Year effect (*T*-test- $p \leq 0.05$ ) was statistically non-significant for both wheat and maize crops.

### 3.11. Effect of irrigation and nitrogen rates on economic and marginal analysis

Economic analysis is considered the ultimate yardstick to judge a treatment effectively in terms of economic analysis and to recommend a specific technology. The economic analysis was performed for both growing seasons, and the average values are presented in Table 9. We found that the value cost ratio (VCR) (which is equal to net return divided by total expenditure) was highest in I2N3 (1.64 and 2.04 in wheat and maize, respectively), which was similar to I3N3 (1.64 and 2 in wheat and maize, respectively). However, the lowest VCR values (1.21 in wheat, 1.24 in maize) were noticed in the I1N1 treatment. The VCR trend in descending order under the interactive effect of irrigation and nitrogen was I2N3>I3N3>I2N2>I3N2 in wheat crop and was I2N3>I3N3>I3N2>I2N2 in maize. The marginal net benefit was highest under I3N3 in wheat (464.1) and maize (917.6), followed by 420.4 under I2N3 in wheat and 870.9 under I3N2 in maize. However, the minimum values of marginal net benefit were 51.9 and 264.3 in wheat and maize, respectively. A similar trend for marginal rate of return was noticed in wheat, such that the greater value (389%) was found in 13N3, followed by I2N3 (371.3%) and I2N2. However, in the maize crop, the highest value (779.5%) of marginal rate of return was noticed in the I2N3 treatment, followed by 772.2% in the I3N3 treatment. Similar to marginal net benefit, the lowest values for marginal rate of return were also noticed in the I1N2 treatment, which were 40.2 and 246.8% in wheat and maize, respectively. Singh et al. [14] developed a relationship between irrigation and yield and stated that proper irrigation scheduling enhances yield, but excessive irrigation could not be cost-effective, as it decreased yield and water use efficiency. Similarly, Si et al. [70] found that optimum nitrogen application has the highest mean return as compared to other nitrogen levels. We noticed that I2 could be a suitable irrigation level in the wheat-maize cropping system without significant economic loss. Apart from irrigation, nitrogen application at the N3 level performed well, and N2 could also be a suitable level if farmers do not have sufficient resources.

**Table 9.**
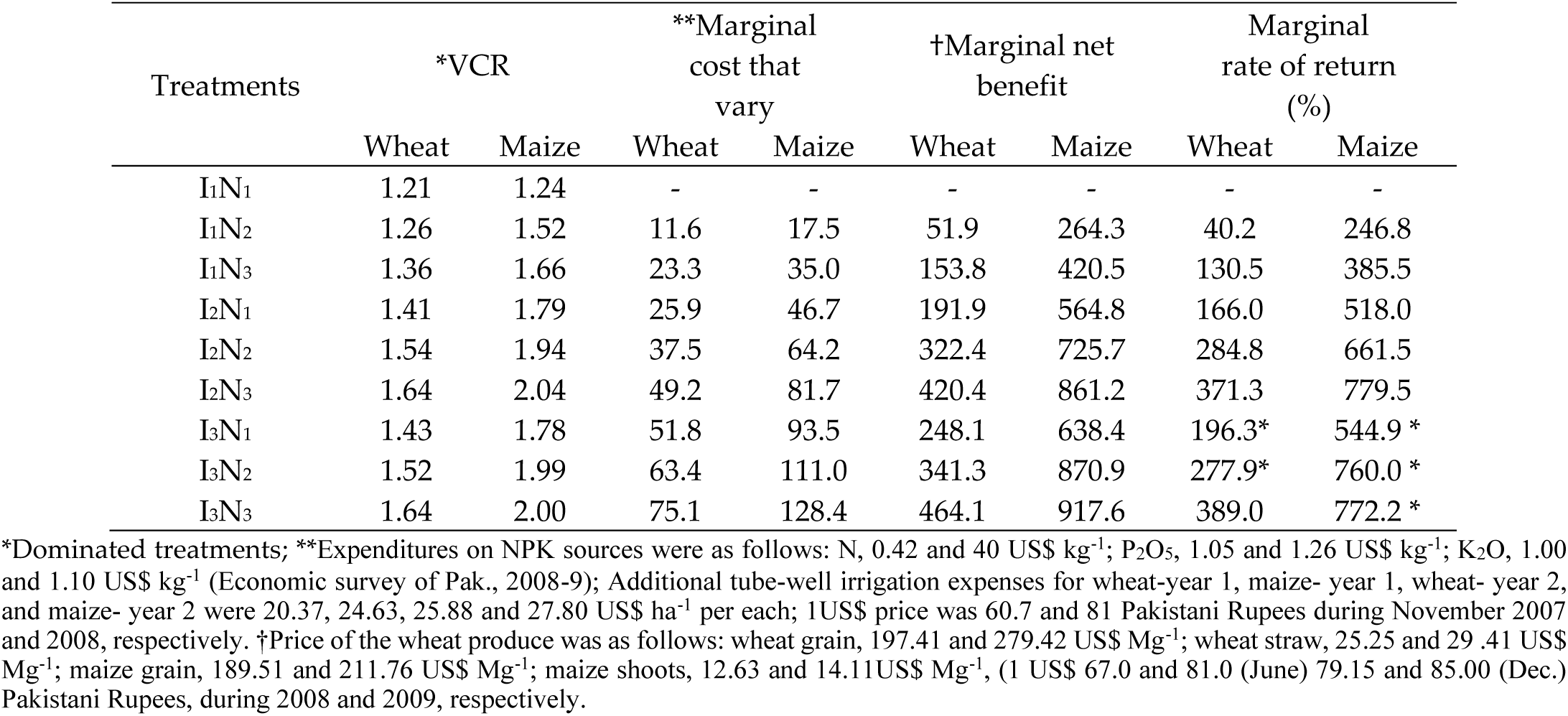
Effect of irrigation and nitrogen rates on economic and marginal analysis of wheat and maize.

| Treatments | *VCR |  | **Marginal cost that vary |  | †Marginal net benefit |  | Marginal rate of return (%) |  |
| --- | --- | --- | --- | --- | --- | --- | --- | --- |
|  | Wheat | Maize | Wheat | Maize | Wheat | Maize | Wheat | Maize |
| I1N1 | 1.21 | 1.24 | - | - | - | - | - | - |
| I1N2 | 1.26 | 1.52 | 11.6 | 17.5 | 51.9 | 264.3 | 40.2 | 246.8 |
| I1N3 | 1.36 | 1.66 | 23.3 | 35.0 | 153.8 | 420.5 | 130.5 | 385.5 |
| I2N1 | 1.41 | 1.79 | 25.9 | 46.7 | 191.9 | 564.8 | 166.0 | 518.0 |
| I2N2 | 1.54 | 1.94 | 37.5 | 64.2 | 322.4 | 725.7 | 284.8 | 661.5 |
| I2N3 | 1.64 | 2.04 | 49.2 | 81.7 | 420.4 | 861.2 | 371.3 | 779.5 |
| I3N1 | 1.43 | 1.78 | 51.8 | 93.5 | 248.1 | 638.4 | 196.3* | 544.9 * |
| I3N2 | 1.52 | 1.99 | 63.4 | 111.0 | 341.3 | 870.9 | 277.9* | 760.0 * |
| I3N3 | 1.64 | 2.00 | 75.1 | 128.4 | 464.1 | 917.6 | 389.0 | 772.2 * |

\*Dominated treatments; \*\*Expenditures on NPK sources were as follows: N, 0.42 and 40 US\$ kg<sup>-1</sup>; P<sub>2</sub>O<sub>5</sub>, 1.05 and 1.26 US\$ kg<sup>-1</sup>; K<sub>2</sub>O, 1.00 and 1.10 US\$ kg<sup>-1</sup> (Economic survey of Pak., 2008-9); Additional tube-well irrigation expenses for wheat-year 1, maize- year 1, wheat- year 2, and maize- year 2 were 20.37, 24.63, 25.88 and 27.80 US\$ ha<sup>-1</sup> per each; 1US\$ price was 60.7 and 81 Pakistani Rupees during November 2007 and 2008, respectively. †Price of the wheat produce was as follows: wheat grain, 197.41 and 279.42 US\$ Mg<sup>-1</sup>; wheat straw, 25.25 and 29.41 US\$ Mg<sup>-1</sup>; maize grain, 189.51 and 211.76 US\$ Mg<sup>-1</sup>; maize shoots, 12.63 and 14.11US\$ Mg<sup>-1</sup>, (1 US\$ 67.0 and 81.0 (June) 79.15 and 85.00 (Dec.) Pakistani Rupees, during 2008 and 2009, respectively.

## 4. Conclusions

Optimum fertilizer N rates applied with efficient irrigation scheduling are important to achieve optimum yield with minimum environmental concerns. Irrigation and fertilizer interactions applied at sub-optimal, optimum, and supra-optimum levels were studied in a wheat-corn rotation for a two-year field experiment. The results from two consecutive years of field experiments based on various irrigation and nitrogen fertilization levels highlighted that optimum irrigation (I2), even at a high nitrogen level (N3), had positive effects on growth, yield, WUEi, and economic profit in the wheat-maize cropping system, with reduced NO_3_-N leaching losses. High nitrogen levels under heavy irrigation, however, led to nitrogen surpluses, which cause soil health and environmental risks in the form of the highest NO_3_-N leaching losses. Moreover, high irrigation water (I3) significantly reduced WUEi, and low irrigation (I1) caused a significant reduction in crop yield, thereby reducing profit from both wheat and maize crops. Thus, a combination of I2 irrigation level with N3 level could be the best irrigation and nitrogen application pattern in the wheat-maize cropping system. The results from this study provide a scientific basis for wheat and maize production with efficient resource utilization through manipulation of various irrigation and nitrogen levels. However, further studies are needed to consider early sowing of the corn crop to reduce significant NO3-N leaching losses during the rainy fallow period.

## Supplementary Materials

The following supporting information can be downloaded at: www.mdpi.com/xxx/s1, Table S1 title: Detail of management operations; Table S2 title: Parameters of water retention curve measured using RETC-fit software applying dual porosity - fit of retention (Durner model), used for drainage calculation using HYDRUS-1D; Table S3 title: Soil sampling and leachates collection schedule for NO_3_-N; Table S4 title: Effect of irrigation and N rates on WUE (kg ha^−1^ mm^−1^) of wheat and maize; and Table S5 title: Statistics of root weight density and root length density.

## Author Contributions

Conceptualization: Muhammad Tahir, Anwar ul Hassan, and David Mulla; methodology, software, validation, and formal analysis: Muhammad Tahir, David Mulla, and Saliha Maqbool; investigation, resources, data curation: Saliha Maqbool and Muhammad Tahir; Writing—original draft preparation: Muhammad Tahir, Saliha Maqbool, and Muhammad Zain; Writing—review and editing, Muhammad Zain, and Muhammad Adeel; visualization, supervision, and project administration: Muhammad Tahir and Anwar Ul Hassan; funding acquisition, Muhammad Tahir, and Anwar Ul Hassan. All authors have read and agreed to the published version of the manuscript.”

## Data Availability Statement

The original contributions presented in this study are included in the article/supplementary material. Further inquiries can be directed to the corresponding author(s).

## Acknowledgments

Authors appreciate Dr. Abdul Ghaffar Niazi’s help in collecting some field data.

