## Supplementary material for "Optimizing the fertilizer N rates at different irrigation levels for optimum yield of wheat and corn at reduced nitrate leaching losses": Supplimentry_data

**Table S1** Detail of management operations

| Management operation | Wheat, 2007-8 | Wheat, 2008-09 | Maize, 2008 | Maize 2008 |
| --- | --- | --- | --- | --- |
| Sowing time | Nov., 30 | Dec., 15 | Aug., 20 | Sept., 1 |
| Variety | AS-2002 | AS-2002 | Pioneer-3062 | Pioneer-3062 |
| Sowing method | hand drill | hand drill | Dibbler | Dibbler |
| Plot size | -----------------------------6.7 m x 13.3 m------------------------------ | | | |
| Plant x Plant (cm) | --------------7 cm----------- | | --------------22.5 cm---------- | |
| Row x Row(cm) | --------------22 cm---------- | | --------------67 cm----------- | |
| Irrigation scheduling |  | |  | |
| I1 (DAS) | --------30, 50, 80 & 110----- | | ------10, 25, 46, 67 & 92------ | |
| I_2_ | ----30, 50, 70, 90 & 110----- | | ----10, 25, 40, 55, 65, 75, 92--- | |
| I_3_ | ---30, 50, 65, 80, 95 & 115-- | | --10, 25, 35, 45, 55, 65, 75, 85, 92-- | |
| Weedicide Type & time  Amount | ---Isoproturon 50 WP (55DAS)--- ------1 kg a.i. ha^-1^ --------- | | -----Atrazine 38SC (30 Das)------ --------0.774 kg a.i. ha^-1^ --------- | |
| Harvesting | Apr., 19 | Apr., 28 | Dec., 12 | Dec., 24 |

DAS, days after sowing of crop; Time between the harvest of preceding wheat crop and succeeding maize crop was fallow period. First irrigation to wheat was 100 mm, while all other irrigations consisted of 75 mm.

**Table S2.** Parameters of water retention curve measured using RETC-fit software applying dual porosity - fit of retention ([*Durner*](mk:@MSITStore:C:\Program%20Files\PC-Progress\HYDRUS-1D%204.xx\Hydrus1D.chm::/HYDRUS1D/References.htm) model), used for drainage calculation using HYDRUS-1D

| Depth  (cm) | *θ*_r_¶  (-) | α_m_  (cm^-1^) | n_m_ | α_im_  (cm^-1^) | ω_im_ | n_im_ | θ_FC_  (-) | θ_PWP_  (-) | θ_AWC_  (-) | SSQ (10^-4^) | r^2^ |
| --- | --- | --- | --- | --- | --- | --- | --- | --- | --- | --- | --- |
| **0-35** | 0.042 | 0.045 | 1.56 | 0.020 | 0.57 | 1.35 | 0.261 | 0.118 | 0.143 | 5.54 | 0.99 |
| **35-70** | 0.038 | 0.055 | 1.63 | 0.030 | 0.62 | 1.42 | 0.253 | 0.117 | 0.136 | 9.43 | 0.98 |
| **70-110** | 0.039 | 0.057 | 1.56 | 0.026 | 0.52 | 1.27 | 0.256 | 0.116 | 0.140 | 2.57 | 0.97 |

¶ Data are average of three repeats

**Modeling WRC according to Durner model**

Table-S2 depicts water retention capacity and the Durner parameters of water retention curve obtained by applying dual porosity model using RETC-fit software. It is obvious from the r^2^ and residual sum of square values that dual porosity model fitted well to the observed data. The highest r^2^ value was observed in case of D_1_ i.e. 0.99 while minimum in case of D_3_, i.e. 0.97. Similar to our good fit of retention curve, so many scientists has used dual porosity model preferably under field condition due to non-uniform water flow (Kohne *et al.*, 2006; Gerke and van Genuchten, 1993a).

**Table S3.** Soil sampling and leachates collection schedule for NO_3_-N

| **Crop with sampling #** | **Days after sowing** | **Soil Sampling** | | | **Leachate collection** | | |
| --- | --- | --- | --- | --- | --- | --- | --- |
|  |  | No. of Irrigations up to sampling/ leachate collection | | | | | |
| Wheat |  | I_1_ | I_2_ | I_3_ | I_1_ | I_2_ | I_3_ |
| 1 | 60 | 2 | 2 | 2 | 1 | 1 | 1 |
| 2 | 90 | 3 | 4 | 4 | 2 | 2 | 2 |
| 3 | harvest | 4 | 5 | 6 | 3 | 4 | 4 |
| 4 | - | - | - | - | 4 | 5 | 6 |
| Fallow | End period (last week of fallow period) | | | | | | |
| Maize |  | | | | | | |
| 1 | 30 | 2 | 2 | 2 | 1 | 1 | 1 |
| 2 | 60 | 3 | 4 | 5 | 2 | 2 | 2 |
| 3 | harvest | 5 | 7 | 9 | 3 | 5 | 5 |

**Table S4.** Effect of irrigation and N rates on WUE (kg ha^-1^ mm^-1^) of wheat and maize

|  | Wheat-2007-08 | Wheat-2008-9 | Maize-2008 | Maize-2009 |
| --- | --- | --- | --- | --- |
| I_1_* | 1.14 | 1.08 | 1.56 b | 1.49 b |
| I_2_ | 1.16 | 1.09 | 1.78 a | 1.83 a |
| I_3_ | 1.15 | 1.14 | 1.84 a | 1.93 a |
| N_1_ | 1.16 | 1.10 | 1.65 | 1.67 |
| N_2_ | 1.16 | 1.08 | 1.77 | 1.78 |
| N_3_ | 1.14 | 1.12 | 1.76 | 1.80 |
| I_1_N_1_ | 1.13 | 1.03 | 1.36 b | 1.28 c |
| - N_2_ | 1.13 | 1.09 | 1.63 ab | 1.51 bc |
| - N_3_ | 1.17 | 1.12 | 1.69 ab | 1.67 ab |
| I_2_N_1_ | 1.19 | 1.05 | 1.75 a | 1.84 ab |
| - N_2_ | 1.16 | 1.07 | 1.80 a | 1.80 ab |
| - N_3_ | 1.13 | 1.14 | 1.79 a | 1.86 ab |
| I_3_N_1_ | 1.15 | 1.23 | 1.85 a | 1.89 a |
| N_2_ | 1.18 | 1.08 | 1.89 a | 2.02 a |
| - N_3_ | 1.13 | 1.11 | 1.79 a | 1.88 ab |
| LSD (p≤ 0.05) I | NS | NS | 0.18 | 0.32 |
| N | NS | NS | NS | NS |
| I x N | NS | NS | 0.37,0.35 | 0.25,0.38 |

Effect of irrigation and nitrogen rates on WUE (kg of grain yield ha^-1^ per mm of ET_c_) of wheat and maize. Data (Table- S4) reveal that both irrigation and nitrogen application had statistically non significant effect on water use efficiency (WUE) of wheat crop during both years as do their interactive effect. However, WUE of maize crop was significantly affected by irrigation levels during both years as do their interactive effect. An increase of 18.2% (yr-1) and 29.8% (yr-2) in WUE of maize was observed with I_3_ treatment over the I_1_. Treatment combination “I_3_N_2_” showed maximum water use efficiency, i.e. 1.89 and 2.02 kg ha^-1^ mm^-1^, during yr-1 and yr-2, respectively.

**Table S5** Statistics of root weight density and root length density

| **LSD (**p≤**0.05)** | **Wheat-1** | **Wheat-2** | **Maize-1** | **Maize-2** |
| --- | --- | --- | --- | --- |
| **RWD** | | | | |
| I | 0.13 | 0.18 | 0.18 | 0.11 |
| N | 0.12 | 0.09 | 0.10 | 0.08 |
| IxN* | 0.20, 0.21 | 0.15, 0.22 | 0.17, 0.22 | 0.14, 0.16 |
| **RLD** | | | | |
| I | 0.03 | 0.50 | 1.29 | 0.70 |
| N | 0.29 | 0.26 | 0.71 | 0.61 |
| IxN | 0.50, 0.41 | 0.45, 0.41 | 1.24, 1.63 | 1.05, 1.09 |
| Note: year effect was statistically non-significant for both crops | | | | |

* 1^st^ LSD value is for same levels of irrigation, while 2^nd^ for different levels of irrigation
